# Beyond mean fitness: demographic stochasticity and resilience matter at tree species climatic edges

**DOI:** 10.1101/2022.08.22.504487

**Authors:** Arnaud Guyennon, Björn Reineking, Roberto Salguero-Gomez, Jonas Dahlgren, Aleksi Lehtonen, Sophia Ratcliffe, Paloma Ruiz-Benito, Miguel A. Zavala, Georges Kunstler

## Abstract

**Aim:** Linking local population dynamics and species distributions is critical to predicting the impacts of climate change. While many studies focus on the mean fitness of populations, theory shows that species distributions can be shaped by demographic stochasticity or population resilience. Here we examine how mean fitness (measured by invasion rate), demographic stochasticity, and resilience (measured by the ability to recover from disturbance) constrain populations at the edges compared to the climatic center.

**Location:** Europe: Spain, France, Germany, Finland, and Sweden.

**Period:** Forest inventory data used for fitting the models cover the period from 1985 to 2013.

**Major taxa:** Dominant European tree species; Angiosperms and Gymnosperms.

**Methods:** We developed dynamic population models covering the entire life cycle of 25 European tree species with climatically dependent recruitment models fitted to forest inventory data. We then ran simulations using integral projection and individual-based models to test how invasion rates, risk of stochastic extinction, and ability to recover from stochastic disturbances differ between the center and edges of species’ climatic niches.

**Results:** Results varied among species, but in general, demographic constraints were stronger at warm edges and for species in harsher climates. Conversely, recovery was more limiting at cold edges. In addition, we found that for several species, constraints at the edges were due to demographic stochasticity and recovery capacity rather than mean fitness.

**Main conclusion:** Our results highlight that mean fitness is not the only mechanism at play at the edges; demographic stochasticity and population capacity to recover also matter for European tree species. To understand how climate change will drive species range shifts, future studies will need to analyse the interplay between population mean growth rate and stochastic demographic processes as well as disturbances.

## 1 Introduction

Given the magnitude of the projected climate changes, the distribution of tree species across Europe is likely to change significantly (Cheaib et al., 2012). Understanding how local population dynamics control tree species range limits is crucial to predict range shifts (Schurr et al., 2012). However, we still have a very crude understanding of this relationship.

The Hutchinsonian niche concept states that species range limits occur in the environmental conditions where population performance does not allow them to persist (Godsoe, Jankowski, Holt, & Gravel, 2017; Hutchinson, 1978). This concept is at the core of a long-standing postulate in biogeography known as the ‘abundant centre hypothesis’ (Brown, 1984; Pironon et al., 2017), which proposes that local population performance decreases towards the range limits. Although species range limits could also be influenced by other processes, such as dispersal and non-equilibrium dynamics (Holt, Keitt, Lewis, Maurer, & Taper, 2005), the focus on the local population dynamics level proposed by this hypothesis has guided empirical studies on range limits. Most of these studies have focused on the importance of a decline of tree population growth rate towards the species range limits (Csergo et al., 2017; Le Squin, Boulangeat, & Gravel, 2021; Purves, 2009). However, theoretical studies have demonstrated that the links between species range limits and local population dynamics could be more complex than just an effect on mean population growth rate (Holt et al., 2005; Sexton, McIntyre, Angert, & Rice, 2009).

Holt et al. (2005) proposed three mechanisms that could lead to stable range limits. The first mechanism is based on the classical idea that species are present where their mean population growth rate allows their presence to be maintained. Previous studies generally used density-independent models and were thus estimating mean finite population growth rate (Csergo et al., 2017). However, for populations with strong density-dependence, such as trees, invasion rate is more appropriate than population growth rate (Le Squin et al., 2021; Pagel et al., 2020; Purves, 2009). The second mechanism, demographic stochasticity, describes the random fluctuations in population size due to probabilistic discrete events of individual tree recruitment and death (quantified by the demographic variance, see Melbourne, 2012, for an in-depth definition), which might ultimately result in local extinction (Grimm & Wissel, 2004). Extinction risk increases when demographic variance increases or when the number of individuals decreases (Engen, Sæther, & Møller, 2001). The third mechanism, environmental stochasticity, assumes that temporal variations in extrinsic environmental conditions, such as climatic or disturbances, may affect population persistence and thus species distribution (Holt et al., 2005; Ovaskainen & Meerson, 2010). In forest ecosystems, the ability of the population to recover from external disturbance is critical (Seidl et al., 2017).

When exploring changes in local population dynamics at species’ range limits, it is, therefore, crucial to analyze these mechanisms jointly. Indeed, demographic and environmental stochasticity remain largely underexplored, yet could explain why populations experience local extinctions at their climatic range limits, even if their population growth rate is positive (Holt et al., 2005). In addition, these three mechanisms are not mutually exclusive, but their respective roles are likely to vary between limits with different physiological constraints. Indeed, Kunstler et al. (2021) recently showed that constraints on tree demographic rates were different between drought- and cold-limited climatic limits, as tolerance to different abiotic stresses requires different adaptive strategies. Finally, even if species were adapted to their environment, the response of local population dynamics may be expected to depend on the climatic optimum of the species, with stronger responses at range limits under more extreme climatic conditions due to the disproportionate effects of climatic extremes (Kunstler et al., 2021).

Recently, several studies have assessed how population dynamics drive tree species distributions using National Forest Inventories (hereafter NFIs) (Kunstler et al., 2021; Le Squin et al., 2021; Purves, 2009; Schultz et al., 2022; Thuiller et al., 2014; Yang et al., 2022). However, to our knowledge, there have been no systematic tests of the respective roles of demographic and environmental stochasticity for range limits of tree species (but see Pagel et al., 2020, for shrub response to fire disturbance in South Africa), probably because most studies either ignored recruitment or assumed it was independent of climate (Kunstler et al., 2021; Le Squin et al., 2021, but see Purves *et al*. 2009). Recruitment, however, is a key stage of the life cycle to properly explore the role of stochastic processes (Grubb, 1977; Holt et al., 2005).

Here, we assessed the relative importance of the three mechanisms presented above on the continental distributions of 25 European tree species (9 gymnosperms and 16 angiosperms). We extended an integral projection model (IPM) recently developed for European tree species (Kunstler et al., 2021) by adding species-specific climate- and density-dependent recruitment models. The IPMs developed here describe the full life cycle of each species. As such they allowed us to estimate metrics of population performance representative of the three mechanisms proposed by Holt et al. (2005) and then test how they differ between the centre and the edges of the species climatic niches (see Fig. 1 for an overview of the metrics and the tests). For each species we differentiated a ‘hot and dry’ and a ‘cold and wet’ climatic edge. More specifically, we tested the following hypotheses: (H1) Mean population performance, measured by the invasion rate, decreases at the edge relative to the center (Brown, 1984). (H2) The risk of stochastic extinction increases at the edge relative to the center because of a higher demographic stochasticity and/or a smaller tree density at equilibrium (Holt et al., 2005). (H3) The ability to recover from stochastic disturbances decreases at the edge compared to the center. We also tested whether the role of the three mechanisms (H1 to H3) differs between the hot and dry edge *vs*. the cold and wet edge (H4). Finally, we tested whether the strength of limitation at the edge is stronger for species’ edges in the extremes of European climate (hot edges of hot-distributed species and cold edges of cold-distributed species, H5).

**Figure 1:**
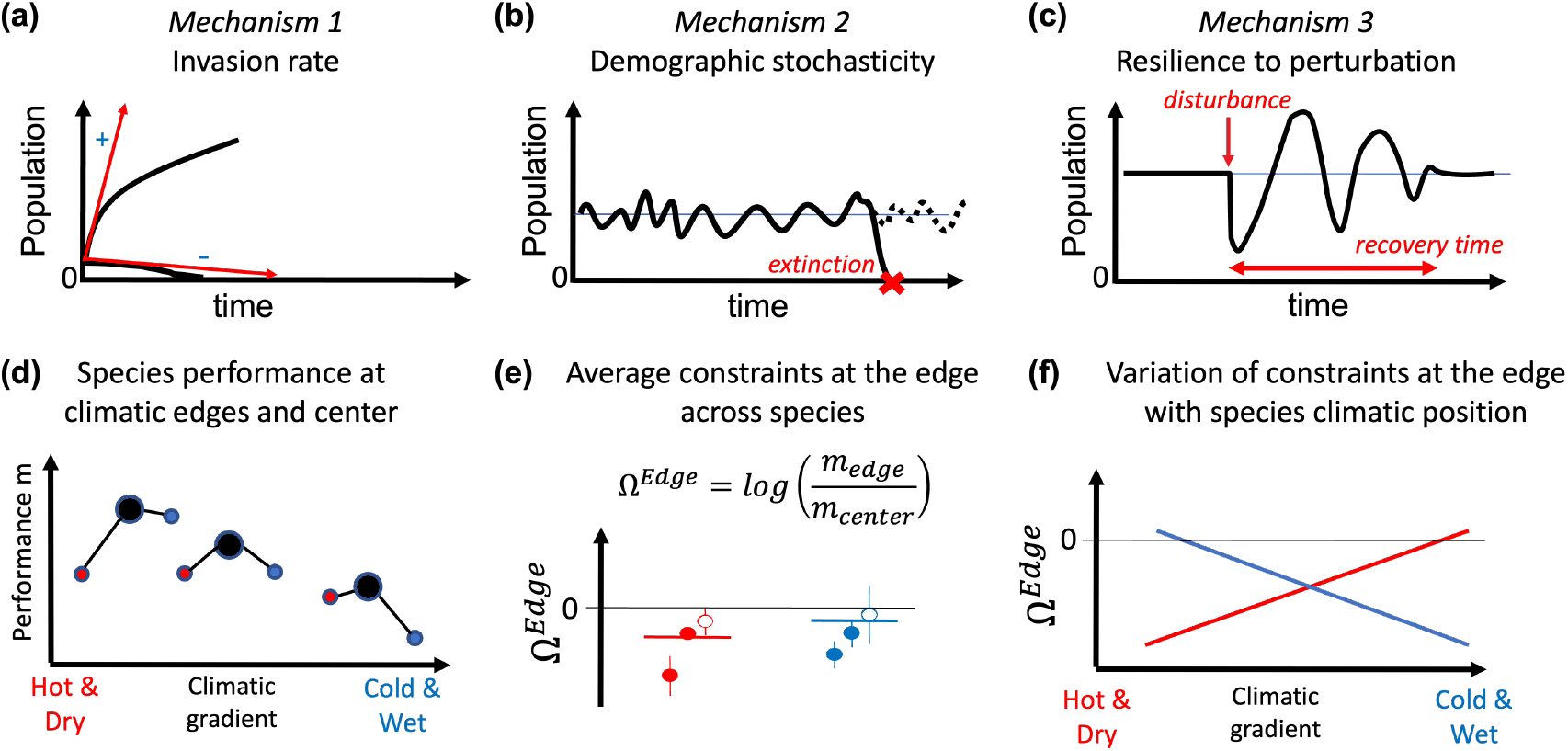
Conceptual figure illustrating the three groups of mechanisms that could limit species distribution at their edges proposed by Holt *et al*. (2005) (a, b, and c), and the approach to test their responses at the hot and dry or the cold and wet edges, and their variation depending on species climatic center (d, e, and f). (a) Mean fitness is estimated by the invasion rate as the population’s ability to grow when rare (black lines represent two different population trajectories of invaders and the red arrows their estimated invasion rates), (b) demographic stochasticity is measured as the variability of tree density solely due to stochasticity of vital rates and its effect on the risk of stochastic extinction (lines represents stochastic tree density variations in small populations that results in extinction for the solid line at the red cross), (c) resilience to disturbance is measured as the recovery time of a tree population after disturbance (represented by the red arrow). (d) Values of population performances *m* for three species at their climatic niche center (black circle), hot and dry edge (red circle) and cold and wet edge (blue circle) (value along the x-axis represents their positions on the climatic gradient), (e) index of response at the edge in comparison to the center – Ω^*edge*^ for the three species, filled points represent significant species responses and the horizontal line represents the overall effect allowing to test if there is a general response of Ω across all species, (f) variations of Ω with species’ climatic niche center (i.e. median of their positions along the climatic axis). The three graphics present the expected results according to our hypotheses: population performance decline at the edges which is equivalent to Ω < 0 at each edge, and Ω decrease is stronger at the hot and dry edges of species occurring in hot climate and at the cold and wet edge of species occurring in cold climate. See Materials and Methods for a full description of the metrics, Ω, and the statistical tests.

## 2 Materials and Methods

### 2.1 Forest Inventory and climatic data

#### National Forest Inventory dataset

To fit vital rate functions (growth, survival, and recruitment), we used the European forest inventory data compiled in the FunDivEUROPE project (Baeten et al., 2013). The dataset contains information on individual trees in 91,528 plots across Spain, France, Germany, Sweden and Finland, with records of species identity, diameter at breast height (dbh), and status (alive, dead, harvested) at two surveys. These data allow to both track individual growth and survival and to describe local competition. We focused our analysis on 25 species with *>* 2,000 individuals and *>* 500 plots in the data, to ensure a good coverage of their range and good estimation of their demographic rates. The mean number of individuals and plots per species were respectively 38531 and 5568 and the census time ranged from 5 to 20 years, with a median of 11 years (more details in Table S2 in Supporting Information). The minimum dbh of trees was 10 cm and no data were available on either seed production by conspecific adult trees, or seedling and sapling growth/survival. We thus did not disentangle the different stages leading to the ingrowth of a 10 cm dbh tree (i.e. trees that grew larger than the 10 cm dbh threshold between two surveys).

Survey design varies between countries, but generally plots are circular with variable radii depending on tree size (largest radius ranging from 10 m to 25 m, see protocols in Supporting Information S1). We excluded from the analyses all plots with records of harvesting operations or disturbances between the two surveys, which would otherwise influence our estimation of local competition.

#### Climate variables

Following Kunstler et al. (2021), we used two climatic variables known to control the physiological performance of trees to fit our vital rates functions (see SI 3.1): the sum of degree days above 5.5°*C* (hereafter *sgdd*), and the water availability index (*wai*). *wai* is calculated as 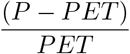 (Sophia Ratcliffe et al., 2017), with *P* the annual precipitation and *PET* the potential evapotranspiration. Daily temperature and *P* were extracted from Moreno and Hasenauer (2016), and *PET* from the Climatic Research Unit data (Harris, Jones, Osborn, & Lister, 2014). Climate variables were averaged over the period covered between two years before the first survey and the year of the last survey, to account for the fact that climate might have a delayed effect on demographic rates.

### 2.2 Integral Projection Model models

An IPM predicts the size distribution, *n*(*z*′, *t* + 1), of a population at time *t* + 1 from its size distribution at time *t, n*(*z, t*), based on a kernel *K*(*z*′, *z*) (with *z* and *z*′ the size at time *t* and *t* + 1) (Easterling, Ellner, & Dixon, 2000). Here, we consider size as the diameter at breast height (dbh). *K*(*z*′, *z*) can be split into the survival and radial growth kernel *P* (*z*′, *z*) and the fecundity kernel *F* (*z*′, *z*), as follows : *K*(*z*′, *z*) = *P* (*z*′, *z*) + *F* (*z*′, *z*). The survival and radial growth (hereafter growth) kernel *P* (*z*′, *z*) is defined as *P* (*z*′, *z*) = *s*(*z*) * *G*(*z*′, *z*), *s* being the survival function and *G* the growth kernel. The fecundity kernel *F* (*z*′, *z*) gives the size distribution of newly recruited trees at time *t* + 1 as a function of the size distribution at time *t*. More details on the IPM are provided in SI 2.1.

Below we describe the fitting of the recruitment, growth and survival functions (more details are provided in SI 2.2 and 2.3). Each of these vital rate functions were fitted separately for each species. The impact of climate on vital rates was modelled through two potential alternative shapes: asymptotic or quadratic polynomial. This allowed us to capture alternative climate responses such as increasing, decreasing, or bell-shaped. To account for uncertainty in the climatic response shape, for each species, we fitted 100 models to 70% of resampled data and selected each time the best climatic response model based on the Akaike information criterion (i.e. lowest AIC; see Burnham & Anderson, 2002). Then, we evaluated the goodness of fit on the remaining 30% of the data (see SI 2.2). In the remaining analysis we used the 100 models to translate the uncertainty in the vital rate functions into the metrics of population dynamics.

#### Recruitment function

We developed a recruitment model that accounted for two main processes: fecundity of the conspecific trees (represented by a power function of the basal area of conspecifics), and the competitive effect of heterospecific and conspecific (represented by an exponential function of their basal area, see SI 2.3). Basal area is the sum of cross-sectional area of trees at 130*cm* above ground per plot area (in *m*^2^*ha*^−1^) and is a good surrogate of tree stand biomass. After thorough exploration of different distributions for the number of recruited trees, we fitted for each species a model with a negative binomial distribution using the approach presented above for the climate response. Because the angle count sampling method used in the German NFI makes recruitment analysis difficult, we excluded this country from the recruitment analysis. We used country-specific intercepts to account for variance due to national specificites (e.g. differences in protocols between NFIs), and an offest for the different number of years between surveys.

Finally, in the IPM, we included a delay in tree recruitment to account for the time it takes for a sapling to reach the minimum dbh, meaning that a newly recruited tree is integrated into the population only after 10 years (see SI 2.4).

#### Growth and survival functions

The radial growth and survival were modelled as functions of dbh, basal area of competitors and climatic conditions (*sgdd* and *wai*) as well as country-specific intercepts (as for recruitment). A normal random plot effect accounting for unexplained variation at the plot level was included in the growth model. No random plot intercept was included in the survival model, because in most plots no individuals died between the surveys, making the estimation of a random plot effect difficult. Growth models were fitted with a log normal distribution. Survival models were fitted with a generalized linear model with a binomial error and a complementary log-log link with an offset representing the number of years between the two surveys to account for variable survey times between plots (Morris, Vesk, & McCarthy, 2013). Models with interactions between the climate variables and both size and competition were also tested, to allow trees to have different climatic response depending on their size or their competitive environment. Equations are presented in SI 2.2, and more details are given in Kunstler et al. (2021).

Harvesting is present in all populations and probably leads to a lower natural mortality rate compared to unmanaged forests. Thus, a fixed harvest rate was added to natural mortality in the kernel *P*. We chose to use the mean annual probability of harvesting over the entire dataset and not to include variability in the harvest rate because we are focusing on the climatic drivers of species distribution and not on the effect of management.

### 2.3 Simulations of population dynamics

We simulated dynamics of discretized size distribution **X**_**t**_ (number of individuals per size classes, corresponding to integration of *n*(*z, t*) over each size class) with a matrix formulation of the IPM as follow:

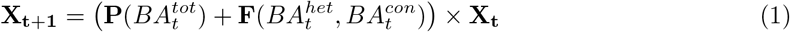

with **P** and **F** the matrices representing the kernel *P* and *F* with the dbh range divided into 700 bins (see SI 2.4 and 2.5 for the numerical integration). Due to the density-dependence of growth, survival, and recruitment rates, the matrix **P** depends on the basal area of competitors at time 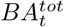, and the matrix **F** on heterospecific and conspecific basal area, respectively 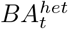 and 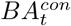.

To explore the effect of demographic stochasticity on the dynamics of small populations, we also developed an individual based model (IBM) based on the same vital rate functions as the IPM (see SI 2.7). In contrast to the IPM, the IBM allows to simulate discrete finite populations which is key to explore demographic stochasticity in small populations. For each species, we ran 100 IBM and IPM simulations using the 100 resampled vital rate functions to represent their uncertainty.

#### Equilibrium

All population metrics, with the exception of invasion rate, were computed starting from equilibrium, because observed tree distributions were highly variable in the forest inventory data. We identified the size distribution at equilibrium *X*_*e*_ for each species and climatic conditions by running simulations with various random initial states until the variations in **X**_**t**_ were negligible. There is no direct analytical solution of the equilibrium for density-dependent IPMs. Still, we checked that our simulations matched the analytical solution for IPMs with a constant transition matrix **P** calculated at the equilibrium basal area (as proposed by Rebarber, Tenhumberg, & Townley, 2012; Townley, Rebarber, & Tenhumberg, 2012, see SI 2.6).

For a small number of species and models, simulations did not reach equilibrium because they predicted a continuous increase in basal area. We discarded models that continued to increase above 200 *m*^2^*ha*^−1^ of basal area at the end of the simulation (the observed maximum basal area in our dataset is 126*m*^2^*ha*^−1^). As simulations work on continuous population abundances, there is no strict extinction. However, there may be very low tree density, which will make the computation of some metrics numerically unstable (recovery from perturbations, for example). In the simulations, we defined a lower limit for basal area of 1*m*^2^*ha*^−1^ (corresponding to one tree of 10 cm in a circle of 10 m diameter) under which populations were not analysed.

If a simulation did not lead to demographic equilibrium (i.e. basal area less than 1*m*^2^*ha*^−1^ or increasing above 200 *m*^2^*ha*^−1^), the simulation was discarded from further analysis. If in a set of simulations less than 50% of the models showed demographic equilibrium, we excluded the full set of simulations (see table S8 in Supplementary Information). In total, only about 9% of the resampled models did not lead to an equilibrium and 5 % of species’ edges were excluded (see SI 2.6).

### 2.4 Population metrics

#### 2.4.1 Invasion rate

Invasion rate was used to evaluate mean fitness. In size-structured populations, the invasion rate is measured by the net reproductive rate, *R*_0_, of a rare invader (Falster, Brännström, Westoby, & Dieckmann, 2017). In our density-dependent IPM, we estimated *R*_0_ by assuming the basal area of the invader was small and had no density-dependent effects on the matrices **F** and **P**. Doing so allowed us to use the same equation as for density-independent IPMs (the dominant eigenvalue of the matrix **F**.(**I** − **P**^−1^), see SI 2.8 and Ellner, Childs, Rees, et al., 2016). As we considered that the invader was rare, we set the conspecific basal area to a low value of 0.1 *m*^2^*ha*^−1^ in *F*. We computed *R*_0_ for two conditions of heterospecific competition: no heterospecific competition (where *BA*^*het*^ = 0), and a high level of heterospecific competition (where *BA*^*het*^ = 60 *m*^2^*ha*^−1^, corresponding to a dense closed forest in our data).

#### 2.4.2 Demographic stochasticity

To evaluate the effect of demographic stochasticity, we derived the time to extinction for finite populations with 250 IBM simulations for each species, climatic condition, and resampled model. We initiated simulations by randomly sampling a finite number of trees from the distribution at equilibrium *X*_*e*_ and for a surface of 100 *m*^2^ and ran the simulation for 1000 years. Following Grimm and Wissel (2004), we extracted the parameter *T*_*m*_ from these simulations, which corresponds to the intrinsic mean time to extinction. While this provides estimates of time to extinction for a very small population that are likely to be much shorter than for large populations in the field, it has the advantage of providing a tool for comparing stochastic extinction between edge and center.

Then, we derived two metrics that drive time to extinction: the density at equilibrium and the demographic variance. Density at equilibrium was computed from long-term simulations, as presented above. We computed the demographic variance from time-series of the total reproductive values (Engen, Lande, Sæther, & Dobson, 2009; Jaffré & Le Galliard, 2016) estimated with long-term IBM simulations (3000 years), on a plot area of 1 *ha* (see SI 2.8).

#### 2.4.3 Population disturbance recovery ability

We used damping time to test a population’s ability to recover from disturbances, and two metrics related to short-term responses. Damping time (i.e. the time to converge to a stable size structure after a disturbance) is independent of the size structure of perturbations (see computation in SI 2.8). This metric, however, does not account for short term transient evolution of the size distribution after a disturbance, as it is computed around equilibrium. Analytical metrics that characterize population transient dynamics can not be used with density dependent models (Capdevila, Stott, Beger, & Salguero-Gómez, 2020). We, thus, used simulations to derive two other metrics: i) *T*_0_ – the time for the first return to equilibrium density (regardless of the tree size distribution); and ii) *T*_*half*_ – the time after which at least half of the initial tree reduction has been permanently recovered. For each species, we disturbed its population at demographic equilibrium by reducing the density of the largest trees (above the diameter 66th percentile) by half and then simulated its dynamics for 1,000 years. We extracted the two metrics from these simulations. For systems that do not present oscillations (i.e. low damping time), these two metrics will be highly correlated.

### 2.5 Response at the edge

#### Niche center and edge definitions

Due to the high correlation between the two climate variables, we defined the climatic position of each species along a single climatic gradient, the first axis of the principal component analysis of the two climatic variables (as Kunstler et al., 2021). For each species, the niche center was the median of the first axis, the hot edge was the 5th percentile and the cold edge was the 95th percentile. To ensure each species’ edges corresponded to valid borders of species distribution, we excluded species and edges where occurrence probability did not decline. Occurrence probability was computed using BIOMOD2 (Thuiller, Lafourcade, Engler, & Araújo, 2009) using presence/absence data from Mauri, Strona, and San-Miguel-Ayanz (2017) (see SI 3.2 and 3.3).

#### 2.5.1 Tests of response at the edge

Using the 100 resampled species-specific IPMs, we predicted the seven metrics at the climatic center (*M*_*center*_), at the hot and dry climatic edge (*M*_*hot*_), and at the cold and wet climatic edge (*M*_*cold*_). We then measured the relative response at the edges as 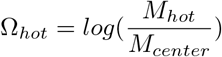 and 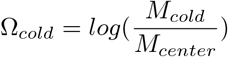. For each metric and edge type we tested whether 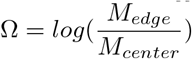 was significantly different from zero (H1 to H4) using a mixed model with edge type effect (hot or cold) as a fixed effect and a random species effect:

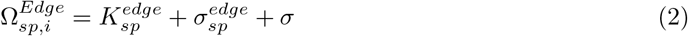

where 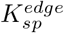 is the edge effect.

To test whether the mean climatic position of the species influenced its response at the edge (H5), we analysed for each metric and edge type the relationship between Ω and the species climatic center conditions. We performed a regression between the value of the median climatic condition and Ω for each edge (taking into account the variance of the 100 resampled models).

As in any comparative analysis, phylogenetic proximity may introduce bias in our analysis, as we may expect similar population metrics between closely related species. That is why we also tested the robustness of the relationships to the phylogenetic proximity of the tree species with a phylogenetic generalized least squares regression (see SI 5, Symonds & Blomberg, 2014).

All analysis were conducted in R cran (R Core Team, 2021), vital rates were estimated using lme4 (Bates, Mächler, Bolker, & Walker, 2014) and glmmTMB (Brooks et al., 2017).

Out of the 27 tree species analyzed, two were fully discarded: *Acer campestre* due to the absence of equilibrium, and *Prunus padus* due to the absence of decline in its prevalence at niche borders.

## 3 Results

### 3.1 Metrics of performance at edges relative to the center

#### Invasion rate

The invasion rate was generally lower at the hot edge than at the center, both in the absence of and at high levels of heterospecific competition, Fig. 2. However, the overall effect across all species was stronger at high levels of heterospecific competition. In the absence of competition, 55% of the studied species had a significantly negative relative response Ω, while at high levels of competition 64% had significantly negative Ω (see Table S7 in SI 4). Relative differences at the cold edge were not significant, with or without competition.

**Figure 2:**
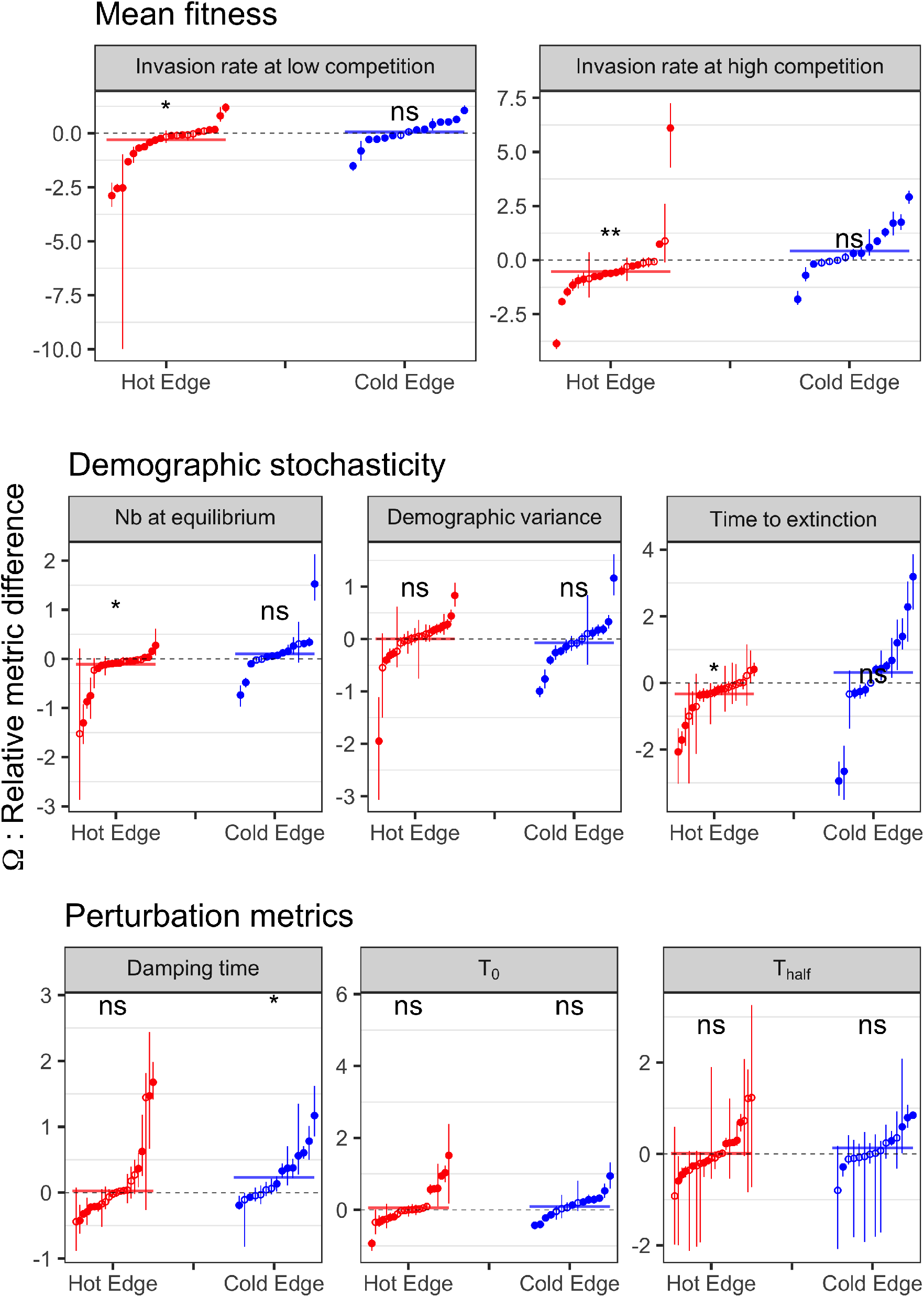
Relative metrics Ω by edge. Each symbol represents a species, error bar is the range of 5 and 95 percentiles. Relative metrics significantly different from 0 (see text) are represented by full circles, otherwise by empty circles. Colored thick horizontal lines represent the edge effect on relative metrics over all species (variable K in equation 2). Significance of relative metrics over all species (see text) is shown with a symbol (ns/*)

#### Demographic stochasticity

The time to extinction was lower at the hot edge compared to the center, with Ω significantly negative for 10 out of 22 species (45%) (Fig. 2). Of the two potential drivers of time to extinction, only Ω of tree density was also significantly less than zero at the hot edge (11 out of 22 species, 50%). Ω of the demographic variance was not significantly positive at the hot edge. No significant effects were detected at the cold edge.

#### Population disturbance recovery ability

Of the three metrics used to study recovery from disturbance, we found a significant effect across all species only for damping time; the damping time tended to be longer at the cold edge compared to the center, indicating slower recovery (8 out of 15 species, 53%) (Fig. 2). There was no difference in damping time between the hot edge and the climatic center. Lastly, we found no differences at either edge type for the time to reach equilibrium density or the time until the perturbation intensity was permanently halved across all species; we found as many species with positive as negative responses.

### 3.2 High species variability in response at the edge

There was a high variability in species response at the edge, with several species showing no effect, or even a higher mean performance at the edge rather than a decrease. This was particularly true for invasion rate without competition, see for example *Abies alba* or *Picea abies* in Fig. 3. Interestingly, for these species, stochastic processes might compensate for the lack of effect on mean fitness at the species level.

**Figure 3:**
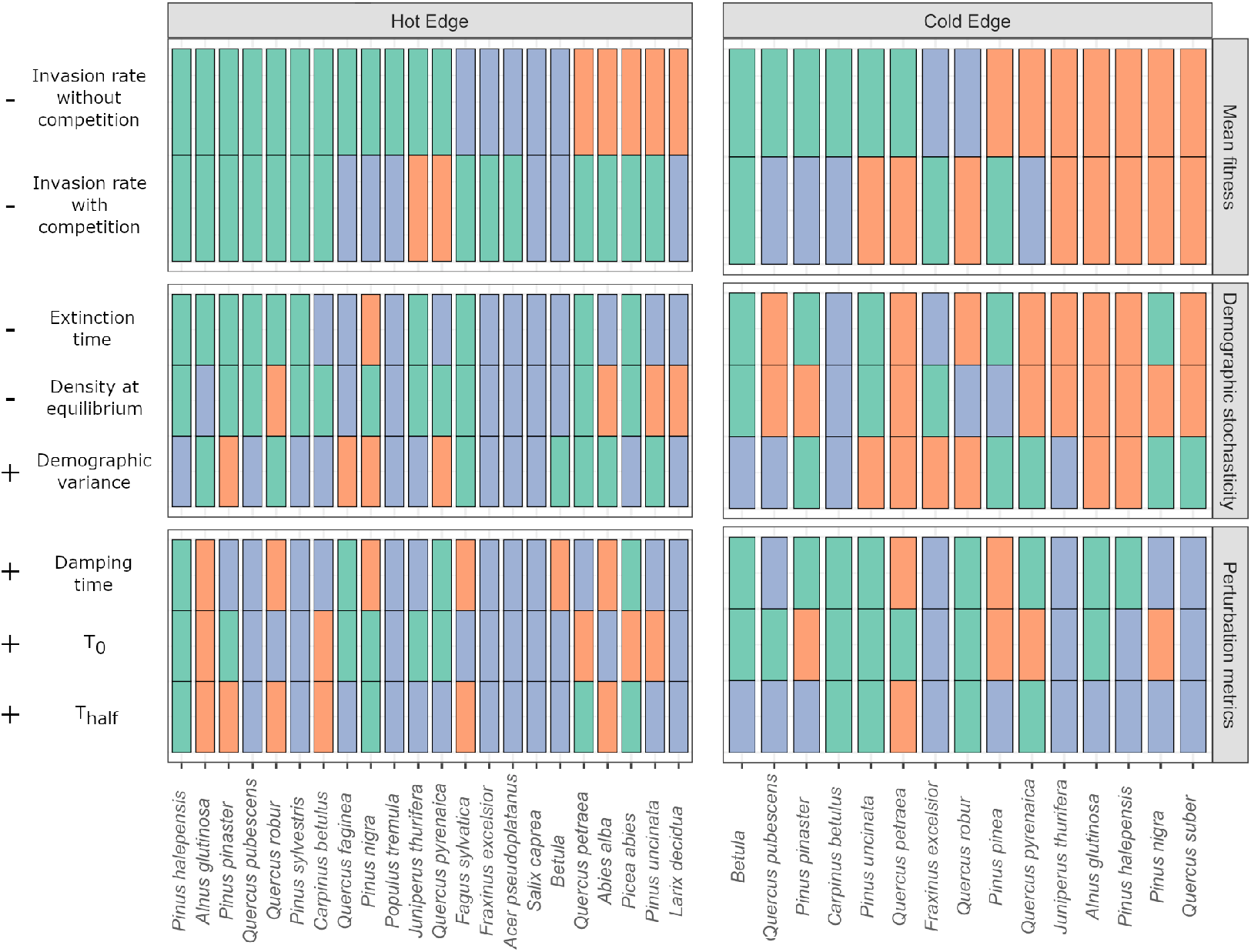
Direction and significance of relative differences of the population performance metrics between edges and climatic center (Ω) for each analysed species at hot and dry edge and the cold and wet edges. Species are ordered from the one showing a significant reduction of invasion rate on the left to the one with opposed response on the right. Green indicates significant constraints on the metric in agreement with the expected direction (expected direction are indicated by – and + signs on the left, decrease or increase in the metric), red indicates a significant effect in opposed direction, and blue indicates a non-significant response.

At the hot edge, among the nine species that did not show a decline of mean fitness or had contrasted mean fitness response (one metric decreased and the other increased), three were constrained by the extinction time (see for example *Juniperus thurifera* or *Quercus petraea* in Fig. 3). At the cold edge, it was the case for three out of ten species (see for example, *Pinus uncinata* or *Pinus nigra* in Fig. 3).

#### 3.2.1 Species responses vary with their climatic center

Part of the variability in species response was related to the position of the climatic center of the species. Several metrics of response at the edge were more severely constrained for species with niche centers in more extreme climates, Fig. 4. At the hot edge, Ω for the invasion rate without competition, tree density, *T*_0_ and *T*_*half*_ were significantly more strongly reduced for species with mean climatic positions in hotter and dryer conditions. At the cold edge, only the invasion rate without competition showed a significant trend, with a stronger reduction in species with a mean climate in colder conditions. These results were robust to the inclusion of phylogenetic structure in the residuals; it only affected the relationship of invasion rates at the cold edge, and damping time at the hot edge (see SI 5).

**Figure 4:**
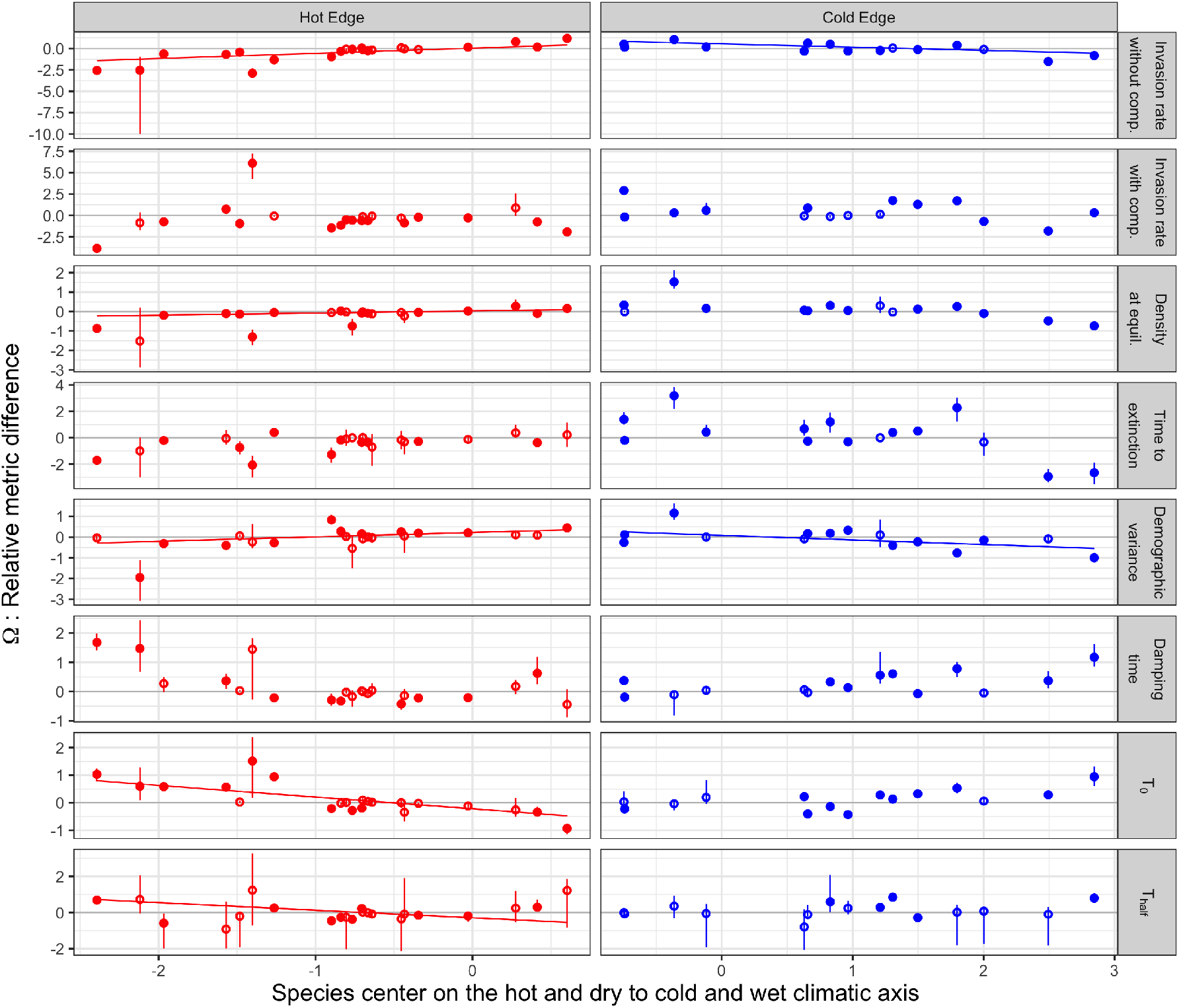
Relative differences of the population performance metrics between edges and climatic center (Ω) along the first principal component axis of species mean climatic conditions. Regression lines are plotted when significant (p-value below 5%). Species relative metrics significantly different from 0 (see text) are represented by full circles, otherwise by empty circles.

## 4 Discussion

Despite considerable variation across species, our results show both a consistent decrease in invasion rate and increase in extinction risk at the hot edge across all species. These patterns were not observed at the cold edge, where only species occurring in extremely cold climates showed a reduction in these two metrics. In contrast, we found a decrease in resilience to perturbation at the cold edge in most species.

### 4.1 Several demographic processes drive species distribution

#### 4.1.1 Invasion rate (H1)

Our results demonstrate a limitation in the invasion rate at the hot edge. This limitation is exacerbated in species that occur in extremely hot and dry climates. These results are consistent with those of a previous study which found that lifespan decreased at the hot edge (Kunstler et al., 2021). The reduced invasion rate at the hot edge is probably the result of this shorter lifespan, but also of the lower recruitment at this edge (observed for species occurring in hot climate, see SI 4.1). Previous studies have proposed that competition could strengthen the limitation on mean fitness at the edge (see Louthan, Doak, & Angert, 2015). Here we found only weak evidence for this, as competition increased the number of species with reduced invasion rate at the edge only from 12 to 14 (see *Quercus petraea* and *Abies alba*, Fig. 3).

We found no clear evidence of a general limitation in invasion rate at the cold edge. Only species distributed at the cold extreme of the gradient showed signs of reduced invasion rate (such as *Betula* and *Pinus uncinata)*. This insight emerged in our new model covering the full life cycle, but not in Kunstler et al. (2021) who did not report a decrease in survival or lifespan for these species. This might be driven by a low recruitment rate at the cold edge for species in extremely cold climates (Figure S15 and S16).

We found that the invasion rate in the absence of competition was more strongly constrained in species from harsh extremes of European climate (hot and dry or cold and wet, hypothesis H5). In contrast, invasion rate was not limited in climatic conditions typical of temperate regions, where productivity is high (Jung et al., 2007).

Direct comparison with previous studies is difficult as they differ in their way of representing species distribution and computing mean fitness (Le Squin et al., 2021; Purves, 2009; Thuiller et al., 2014). However, even if the structure of the model is different from ours, it is interesting that Purves (2009), in a study on East North American tree species, found a significant decrease in invasion rate at the northern edge but not at the southern edge. The fact that fitness decreases occur at the opposite edges for tree species in East North America and in Europe might be related to differences in climatic space between these continents. The European southern edge corresponds to a hot and dry climate, whereas the southern edge of tree species in East North America is not limited by drought (Zhu, Woodall, Monteiro, & Clark, 2015).

#### 4.1.2 Demographic stochasticity (H2)

The mean time to extinction represents an integrative metric of the demographic stochasticity which increases when tree density decreases and demographic variance increases (Ovaskainen & Meerson, 2010). At the hot edge among the 10 species showing a shorter time to extinction, this decline could be related to a change in either demographic variance or tree density or both. This suggests that these two processes could reinforce each other to result in a stronger reduction in the time to extinction. Our results are also interesting in light of the abundant-center hypothesis (Brown, 1984), which postulates a decrease in tree density at the edge of a species range. Indeed, our analysis of tree abundance at long term equilibrium showed that this hypothesis is far from being supported for all edges and species, even with the abundance predicted at equilibrium. This is in agreement with previous large-scale analyses of observed tree abundance Dallas, Decker, and Hastings (2017), Pironon et al. (2017), and Sagarin and Gaines (2002).

#### 4.1.3 Population recovery after disturbances (H3)

Holt et al. (2005) stated that increases in environmental variability can explain range limits despite the absence of a decrease in mean fitness. Here, we explored the role of the time to recovery from disturbances. Disturbance is a key component of environmental variability for tree species. We found an overall significant increase of damping time at the cold edge. This changes in a metric of long term recovery might be connected to the slower tree growth at this edge reported by Kunstler et al. (2021). In contrast, the effect for the short and midterm metrics of population recovery (*T*_0_ and *T*_*half*_) were extremely variable between species (yet, seven species out of 14 showed a longer time to return to the equilibrium density). It is noteworthy that these metrics are extracted from simulations that might lead to a higher variability than the analytical approach used for damping time. At the hot and dry edge, species variability was extremely large. We found evidence of an increase of short and midterm metrics only for species with a climatic center in extreme hot and dry conditions.

A key limitation of our approach on disturbance is that we only explored a single type of abstract disturbance, whereas the real disturbance regime might vary across the species range and play a role in setting distribution limits (Schultz et al., 2022; Senf & Seidl, 2021; Sheil, 2016). In addition, it would be crucial to also explore how interannual variability in climatic conditions, another key component of environmental variability, affects population dynamics. This interannual variability can either have negative, null, or positive effect on the population performance depending on the demographic rates’ response (Le Coeur, Yoccoz, Salguero-Gómez, & Vindenes, 2022). Estimating how natural disturbances and interannual climatic variability might affect tree vital rates and population dynamics at the continental scale remains, however, challenging.

### 4.2 Stronger constraints at the hot edge (H4)

At the hot and dry edges, we found that the invasion rate was constrained and we observed increased stochastic risk of extinction for numerous species. Conversely, constraints at the cold edge were less clear, with an indication of a lower resilience in general and a reduced invasion rate only for species in extreme cold conditions.

These differences might emerge if drought directly results in an increased mortality and higher extinction risk, whereas cold stress could reduce vital rates and population dynamics and thus mechanically increase its response time to disturbances. These differences might also reflect a degree of disequilibrium between the current and potential distribution (Guyennon et al., 2022). Indeed, global warming may lead to an increase in drought pressure at the hot edge (Carnicer et al., 2011) and in contrast a decrease in mortality at the cold edge in Europe (Neumann, Mues, Moreno, Hasenauer, & Seidl, 2017). Such disequilibrium in the southern and northern edges has already been identified in East North America (see Talluto, Boulangeat, Vissault, Thuiller, and Gravel (2017)).

### 4.3 On the challenges of connecting population dynamics and species distribution

It is striking that most studies (including this one) found limited concurrence between mean fitness and species distribution (Kunstler et al., 2021; Le Squin et al., 2021; Purves, 2009; Thuiller et al., 2014). The novelty of our study is that we show that when mean fitness is not constrained at the edge, stochastic processes can play a key role. Yet, there is still a large variability in species responses, with several species having no clear indication of performance constraints or even better performances at edges (*Salix caprea* and *Larix decidua* at the hot edge and*Juniperus thurifera* at the cold edge). Several factors might explain the results for these species. First, we explored species distribution in climate space using only two key climatic variables. Even if these variables discriminate well the distribution of the 25 tree species in Europe (see Fig. S14), species distribution might be influenced by other climatic variables, or other abiotic factors such as soil variables. Secondly, beside environmental space, species distribution can also be analyzed in geographic space (see Pironon et al., 2017). In geographic space, dispersal limitation and decrease in suitable habitat availability can also explain species range limits in a metapopulation framework (Holt & Keitt, 2000). Thirdly, species distributions are not necessarily in equilibrium with current climatic conditions. Highly managed species (such as *Pinus pinaster* or *Picea abies*) can be planted outside their native range. Svenning and Skov (2004) also argued that tree species might still be in the process of slow recolonization since the last glacial age. Here, by initiating our simulations at equilibrium, we effectively removed all legacy effects. Then, as discussed above, climate change could modify constraints at edges (Clark et al., 2021), and explain the difference observed at the hot *vs*. the cold edge. Finally, our ability to capture the complex population dynamics of long-lived organisms such as trees is still limited and might explain the poor match with the distribution. For instance, our models do not consider potential variability in seed production, juvenile growth or survival, which could however also constrain species ranges (Clark et al., 2021). Indeed, we modelled recruitment with a constant lag to account for the time to grow from a seed to a 10 cm of DBH tree across species and climate. This shows that our knowledge on the recruitment phase is still extremely limited. In addition, we explored the role of competition in a relative crude way, considering the competitor effect only through basal area and ignoring the complexity of multispecies interactions. A full exploration of its role would require analysing how the stochastic dynamics of multispecies community constrains species range (Godsoe et al., 2017).

## 5 Conclusion

Our study is one of the first to tease apart several mechanisms that could lead to species range limits using field data across a large set of species at the continental scale. Our results show that the mean fitness may not be the only mechanism at play at the edge; demographic stochasticity and population recovery ability also matter for European tree species. Thus, to understand how climate change will drive species range shifts, we encourage ecologists to analyse the full life cycle of trees and explore how the average population growth rate interacts with stochastic processes and recovery from disturbances in driving species ranges.

## Acknowledgments

This work was funded by the REFORCE - EU FP7 ERA-NET Sumforest 2016 through the call “Sustainable forests for the society of the future”, with the ANR as national funding agency (grant ANR-16-SUMF-0002). GK was funded by the ANR DECLIC (grant ANR-20-CE32-0005-01). GK and BR were funded by BiodivERsA ERA-NET Cofund project FUNPOTENTIAL (ANR funding grant number ANR-20-EBI5-0005-03) and RESONATE H2020 project (grant 101000574). The NFI data synthesis was conducted within the FunDivEUROPE project funded by the European Union’s Seventh Programme (FP7/2007–2013) under grant agreement No. 265171. We thank Gerald Kandler (Forest Research Institute Baden-Wurttemberg) for his help building the German data. We thank the MITECO (“Ministerio para la Transición Ecológica y Reto Demográfico”), the Johann Heinrich von Thunen-Institut, the Natural Resources Institute Finland (LUKE), the Swedish University of Agricultural Sciences, and the French Forest Inventory (IGN) for making NFI data available. GK, SR, and RSG initiated this work in the working group sAPROPOS - ‘Analysis of PROjections of POpulationS’, which was supported by sDiv (Synthesis Centre of the German Centre for Integrative Biodiversity Research - iDiv), funded by the German Research Foundation (FZT 118). MAZ was supported by grant DARE (RTI2018-096884-B-C32, MCINN, Spain). We are grateful to Fabian Roger for providing code to build the species phylogeny.

