## Supplementary Information for "Beyond mean fitness: demographic stochasticity and resilience matter at tree species climatic edges"

**5 Phylogenetic regression along climate** **31**

**References** **33**

### 1 Data description

#### 1.1 Description of the national forest inventories of each country

Table S1 summarises information on survey protocol for each national forest inventory. Below, we give a more detailed description of the protocol for each country.

**Finnish National Forest Inventory** We used data from the eighth NFI (NFI8) of Finland sampled in the period 1985-1986 to 1995. The sample plots are located in a systematic grid across the country of plot clusters in forested areas (Mäkipää & Heikkinen, 2003). In Southern Finland the grid is 16 km by 16 km square, with four plots in each cluster at 400 m intervals, while in Northern Finland the grid is a 24 km by 32 km rectangle with three plots per cluster, at 600 m intervals. These permanent plots were sampled using a variable radius technique with two concentric circular subplots of radius 5.64 m (i.e. 100 $m^2$) for trees with a diameter at breast height (DBH) below 10.5 cm and 9.77 m (i.e. 300 $m^2$) for trees with a DBH above 10.5 cm. The data used in the analysis are provided (with blurred geographic coordinates) in the Dryad Digital Repository https://doi.org/10.5061/dryad.wm37pvmkw (Ratcliffe et al., 2020).

**French National Forest Inventory** The French NFI (IFN 2014) is based on a systematic 1 $km^2$ square grid covering the entire country. A forest is defined in the French NFI as a stand of more than 0.05 ha and wider than 20 m in which crowns from forest trees can reach 5 m and cover more than 10 percent of the area. The whole grid is measured in 5 years and the 5-year sample is divided into five systematic annual sub-samples. Trees are measured in three concentric plots, depending on their circumference at 1.3 m height. Trees with a circumference above 23.5 cm (corresponding to a DBH of 7.5 cm) are measured on a 6 m radius plot; trees with a circumference above 70.5 cm (DBH = 22.4 cm) are measured on a 9 m radius plot; trees with a circumference above 117.5 cm (DBH = 37.4 cm)

| Country | Protocol method | Survey Dates | Plot radius/ minimum DBH | Plot radius/ minimum DBH | Plot radius/ minimum DBH | Plot radius/ minimum DBH |
|---|---|---|---|---|---|---|
| Spain | Variable radius | 1986/1996 1997/2007 | 5 m / 7.5 cm | 10 m / 12.5 cm | 15 m / 22.5 cm | 25 m / 42.4 cm |
| France | Variable radius | Yearly 2005/2011 | 6 m / 7.5 cm | 9 m / 22.5 cm | 15 m / 37.5 cm | |
| Sweden | Variable radius | 2005/2010 2008/2010 | 10 m / 10 cm | | | |
| Finland | Variable radius | 1985/1986 1995 | 5.64 / <10.5 cm | 9.8 m >10.5 | | |
| Germany | Angle count 4 $m^2 ha^{-1}$ | 1986/1990 2001/2002 | | | | |

Table S1: **Description of the protocol of the NFI in each country.**

are measured on a 15 m radius plot; trees with a circumference below 23.5 cm are not measured. For living trees, radial growth over the last five years is measured on short cores. Past 5 years tree mortality is also evaluated.

We used NFI data covering the 2007-2013 period (see https://inventaire-forestier.ign.fr/spip.php?article532).

**German National Forest Inventory** We used information from the first and second German NFI. We used information from the first and second German NFI (https://bwi.info/Download/de/BWI-Basisdaten/ACCESS2003/). The German NFI uses a systematic grid of clusters, sampled during the periods 1986-1990 (undertaken in West Germany only) and 2001-2002. The size of the sample grid is 4 km by 4 km, however, it is reduced in some federal states to either 2.83 km by 2.83 km or 2 km by 2 km. Each cluster is a quadrangle of 150 m in length with a sample plot on each corner (Kandler, 2009). Trees with a DBH above 10 cm in the first inventory and above 7 cm in the second were selected by the angle-count method with a basal area factor (BAF) of 4 $m^2ha^{-1}$ if they are alive or recently dead.

**Spanish National Forest Inventory** We used information from the second and third Spanish NFI (surveyed during the periods 1986-1996 and 1997-2007, respectively) (https://www.miteco.gob.es/en/biodiversidad/servicios/banco-datos-naturaleza/informacion-disponible/ifn2_descargas.aspx and https://www.miteco.gob.es/en/biodiversidad/servicios/banco-datos-naturaleza/informacion-disponible/ifn3_bbdd_descargas.htm.aspx). The Spanish NFI plots are located on a 1 $km^2$ grid over forested regions (Villaescusa & Díaz, 1998; Villanueva, 2004). Spanish NFI plots were sampled using a variable radius technique with four concentric circular subplots of radius 5 m for trees with DBH smaller than 12.4 cm, 10 m for those with DBH between 12.5 and 22.4 cm, 15 m for those with DBH between 22.5 and 42.4 cm and 25 m for those with DBH greater than 42.5 cm.

**Swedish National Forest Inventory** The permanent Swedish inventory uses a regular sampling grid and includes about 4,500 permanent tracts, each surveyed every five years. Plots in the first census were surveyed between 2003 and 2005 and plots in the second census were surveyed between 2008 and 2010. The tracts are rectangular and have different dimensions depending on the location within the country; each tract has between four and eight circular sample plots. All trees with a DBH above 10 cm are sampled in a 10 m radius. The data used in the analysis are provided (with blurred geographic coordinates) in the Dryad Digital Repository https://doi.org/10.5061/dryad.wm37pvmkw (Ratcliffe et al., 2020).

#### 1.2 Summary of observations

Table S2 summarises the number of observations used to fit vital rates.

| Species | Number of trees | Number of dead trees | Number of recruits | Number of plots | Number of plots with death | Number of plots with recruits | Mean Time between surveys | country |
|---|---|---|---|---|---|---|---|---|
| *Abies alba* | 20006 | 444 | 1114 | 3315 | 288 | 580 | 8 | GR/SP/FR |
| *Acer campestre* | 4548 | 106 | 674 | 2084 | 75 | 493 | 6 | SP/GR/FR |
| *Acer pseudoplatanus* | 5027 | 52 | 485 | 2031 | 41 | 317 | 8 | GR/SP/SW/FR |
| *Alnus glutinosa* | 9097 | 354 | 1116 | 1741 | 193 | 428 | 7 | GR/SP/SW/FR |
| *Betula* | 38509 | 1366 | 5599 | 9447 | 999 | 2722 | 6 | GR/SP/FI/SW/FR |
| *Capinus betulus* | 20426 | 299 | 2715 | 5307 | 195 | 1636 | 6 | GR/SW/FR |
| *Fagus sylvatica* | 65775 | 1213 | 3920 | 11052 | 666 | 1820 | 9 | GR/SP/SW/FR |
| *Fraxinus excelsior* | 15439 | 297 | 1507 | 4779 | 222 | 880 | 7 | GR/SP/SW/FR |
| *Juniperus thurifera* | 7293 | 61 | 1092 | 1361 | 22 | 643 | 12 | SP/FR |
| *Larix decidua* | 4139 | 164 | 119 | 969 | 84 | 55 | 8 | GR/SP/FR |
| *Picea abies* | 108211 | 2650 | 9117 | 13884 | 1406 | 3389 | 8 | GR/FI/SP/SW/FR |
| *Pinus halepensis* | 66949 | 2509 | 13128 | 7478 | 683 | 4286 | 11 | SP/FR |
| *Pinus nigra* | 67285 | 2784 | 13814 | 6216 | 470 | 3046 | 11 | SP/GR/FR |
| *Pinus pinaster* | 112497 | 11856 | 13417 | 7837 | 2129 | 3005 | 11 | SP/FR |
| *Pinus pinea* | 20640 | 1238 | 2480 | 2350 | 347 | 735 | 11 | SP/FR |
| *Pinus sylvestris* | 204669 | 7298 | 26465 | 19357 | 2571 | 6441 | 9 | SP/GR/FI/SW/FR |
| *Pinus uncinata* | 14302 | 275 | 1953 | 920 | 117 | 487 | 11 | SP/FR |
| *Populus tremula* | 6660 | 377 | 1016 | 2109 | 250 | 503 | 6 | GR/SP/FI/SW/FR |
| *Prunus padus* | 3145 | 151 | 370 | 1718 | 128 | 293 | 5 | GR/SP/FR |
| *Quercus faginea* | 14031 | 203 | 4111 | 2858 | 90 | 1621 | 11 | SP |
| *Quercus ilex* | 71987 | 1605 | 16824 | 12997 | 750 | 6121 | 11 | SP/FR |
| *Quercus petraea* | 36806 | 899 | 1770 | 7632 | 581 | 815 | 7 | GR/SP/FR |
| *Quercus pubescens* | 26232 | 641 | 2937 | 4879 | 393 | 1582 | 6 | SP/FR |
| *Quercus pyrenaica* | 29056 | 1218 | 7068 | 3189 | 375 | 1782 | 11 | SP/FR |
| *Quercus robur* | 44100 | 1190 | 2489 | 10726 | 819 | 1251 | 7 | SP/GR/FR |
| *Quercus suber* | 19486 | 431 | 1495 | 2655 | 255 | 681 | 11 | SP/FR |
| *Salix caprea* | 4044 | 478 | 1048 | 1453 | 273 | 551 | 5 | SP/SW/FR |

Table S2: **Summary of observations used to fit vital rates by species**. Country codes: GR for Germany, FI for Finland, FR for France, SP for Spain and SW for Sweden. Death and recruits are observed over varying time periods.

#### 2 Presentation of the IPM

##### 2.1 Description of the Integral Projection Model

A size-structured IPM predicts the size distribution, $n(z', t+1)$, of a population at time t+1 from its size distribution at time t, $n(z, t)$, with $z$ the size at t and $z'$ the size at $t+1$, based on the following equation (Ellner, Childs, Rees, et al., 2016):

$$n(z', t+1) = \int_{L}^{U} K(z', z) n(z, t) dz \tag{S1}$$

with $L$ and $U$ being, respectively, the lower and upper observed sizes for integration of the kernel $K$.

The kernel $K(z', z)$ can be split into the survival and growth kernel $P(z', z)$ and the fecundity kernel $F(z', z)$, as follows : $K(z', z) = P(z', z) + F(z', z)$. The fecundity kernel $F(z', z)$ gives the size distribution of newly recruited trees at time $t+1$ as a function of the size distribution at time t. The survival and growth kernel $P(z', z)$ is defined as $P(z', z) = s(z) * G(z', z)$, $s$ being the survival function and $G$ the growth function.

The figure S1 presents the structure and approach of the IPM.

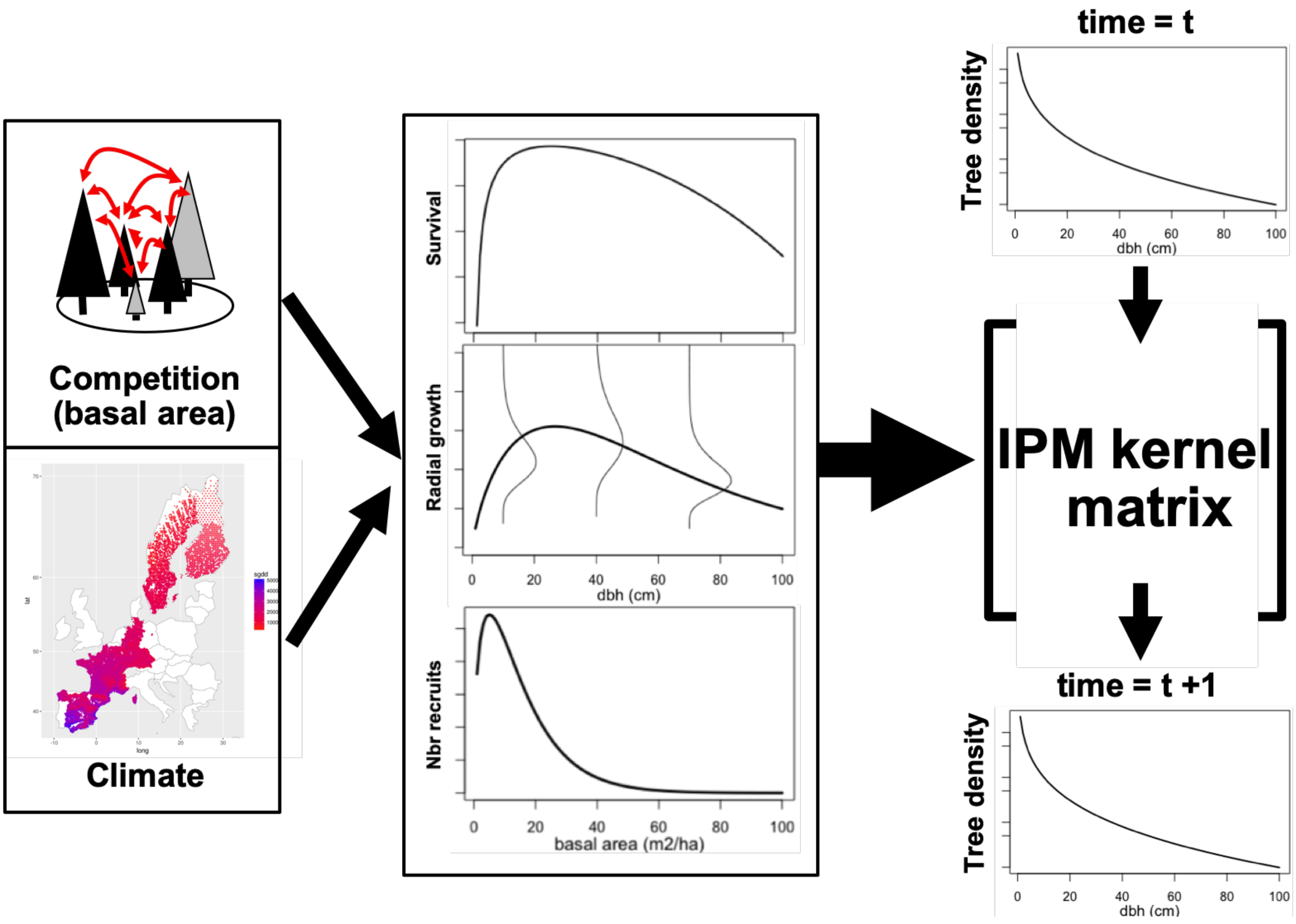

Figure S1: **Schematic representations of the IPM and the vital rate functions**.

##### 2.2 Growth and survival functions

Below, we present the equations of the growth and survival functions that were developed in Kunstler et al. (2021).

##### 2.2.1 Growth function

The growth functions are based on the two following minimal equations:

$$
\begin{aligned}
log(G_{i,p}) =& a_0 + a_{0,p} + a_1\, D_i + a_2\, log(D_i) + \\
& a_3\, BA_i + a_4\, \frac{1}{sgdd_p} + a_5\, \frac{1}{wai_p} + \varepsilon_i
\end{aligned}
\tag{S2}
$$

$$
\begin{aligned}
log(G_{i,p}) =& a_{0,c} + a_{0,p} + a_1\, D_i + a_2\, log(D_i) + a_3\, BA_i + \\
& a_4\, sgdd_p + a_5\, sgdd_p^2 + a_6\, wai_p + a_7\, wai_p^2 + \varepsilon_i
\end{aligned}
\tag{S3}
$$

Where $G_{i,p}$ is the annual diameter growth of tree $i$ in plot $p$, $D_i$ is the dbh of tree $i$, $BA_i$ is the sum of basal area of local competitors of tree $i$ per ha (sum basal area of both conspecific and heterospecific trees in the plot in a single local competition index), $sgdd_p$ is the sum of growing degree days, $wai_p$ is the water aridity index, $a_0$ to $a_7$ are estimated parameters, and $a_{0,p}$ is a normal random plot effect accounting for unexplained variation at the plot level. The intercept $a_{0,c}$ is country-specific to account for differences in sampling protocols between the NFIs (plot size and difference in mean survey time) and $\varepsilon_i$ is the unexplained tree level variability following a normal distribution. For the IPM simulations we used the average of the intercept across countries. We also tested models with interactions between the climatic variables - $1/sgdd_p$ and $1/wai_p$ for model (S2) and $sgdd_p$ and $wai_p$ for model (S3)) - and size ($D_i$ and $log(D_i)$) and the climatic variables and competition. We fitted the models in R-cran separately for each species (R Core Team, 2019) using the 'lmer' function ("lme4" package, Bates, Mächler, Bolker, & Walker, 2015).

##### 2.2.2 Survival function

The survival functions are based on the two following minimal equations:

$$
\begin{aligned}
cloglog(S_{i,p}) =& a_{0,c} + a_1\, D_i + a_2\, log(D_i) + \\
& a_3\, BA_i + a_4\, \frac{1}{sgdd_p} + a_5\, \frac{1}{wai_p} + log(y_p)
\end{aligned}
\tag{S4}
$$

$$
\begin{aligned}
cloglog(S_{i,p}) =& a_{0,c} + a_1\, D_i + a_2\, log(D_i) + a_3\, BA_i + \\
& a_4\, sgdd_p + a_5\, sgdd_p^2 + a_6\, wai_p + a_7\, wai_p^2 + log(y_p)
\end{aligned}
\tag{S5}
$$

Where $S_{i,p}$ is an integer with value 1 if the tree $i$ survived between the two surveys and 0 if the tree died, $y_p$ is the number of years between the two surveys added as an offset. $D_i$ is tree diameter at breast height $i$, $BA_i$ is the sum of basal area of competitors for tree $i$ per ha, and $sgdd_p$ and $wai_p$ are respectively the sum of growing degree days and the water aridity index of the plot $p$. $a_0$ to $a_7$ are fitted parameters. $a_{0,c}$ is a country specific intercept to account for differences of protocol between countries. For the IPM simulations, we used the average of the intercept across countries.

##### 2.2.3 Growth and survival functions response curves

Figures S2 and S3 represent the vital rates (survival and radial growth, respectively) at the different niche positions of the tree (hot edge, center, and cold edge). Survival and radial growth depend on tree diameter; competition was set to null in these figures. The parameters of the 100 refitted models of each species are provided in '.rds' R object in the Zenodo repository, https://zenodo.org/record/6424490 (for instance for *Fagus sylvatica* in the file 'output/fit_sgr_all_Fagus sylvatica.Rds').

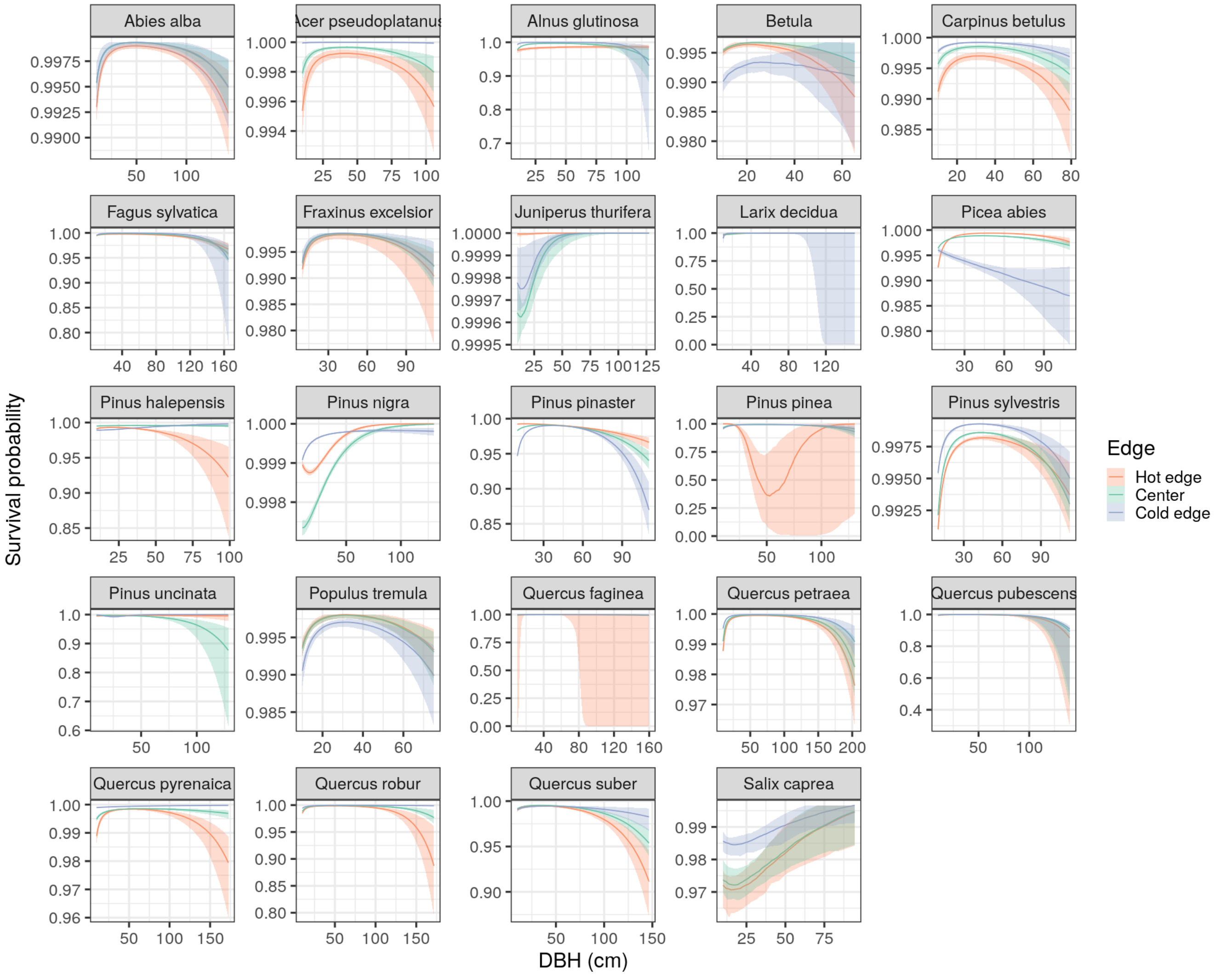

Figure S2: **Survival rates according to tree diameter**. Each color corresponds to a climatic niche position, lines represent the median value between all 100 sub-models, and the shaded areas the interval between the 5th percentile and the 95th percentile.

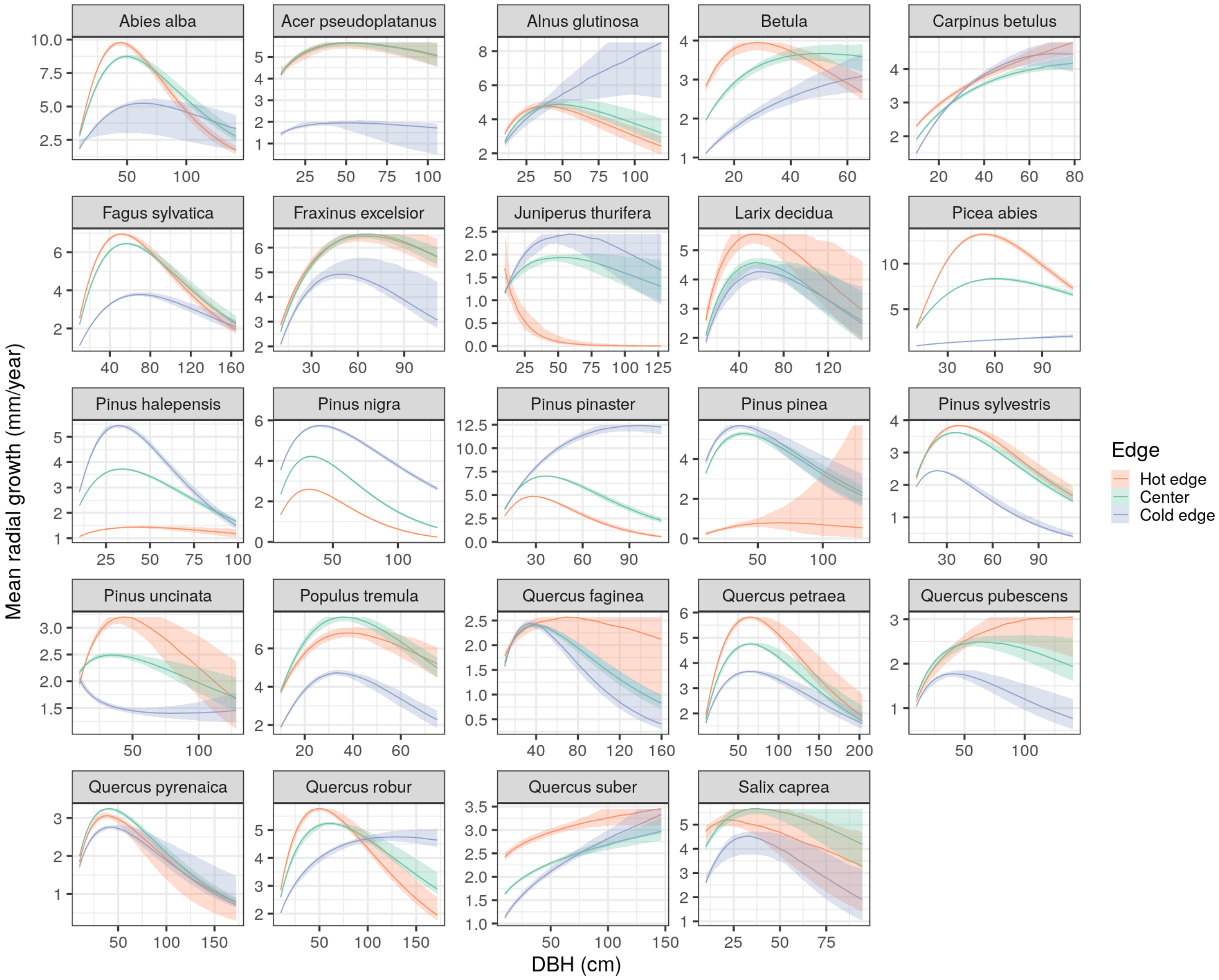

Figure S3: **Radial growth rates according to tree diameter**. Each color corresponds to a climatic niche position, lines represent the median value between all 100 sub-models, and the shaded areas the interval between the 5th percentile and the 95th percentile.

##### 2.2.4 Growth and survival goodness of fit

We cross-validated the 100 growth and survival models fitted on a re-sampling of 70% of data on the remaining 30% by computing the normalised root-mean-square error (NRMSE) for growth and area under the curve (AUC) for the survival model (Figures S4 and S5). Due to the larger amount of data in some areas of the climatic niche, the climatic dependencies estimations may be biased. For each species, the re-sampling was performed to have higher probability of sampling in climatic conditions with fewer data (the climatic gradient is divided into 30 classes and the probability of re-sampling for plots in each class is inversely proportional to the number of plots in that class).

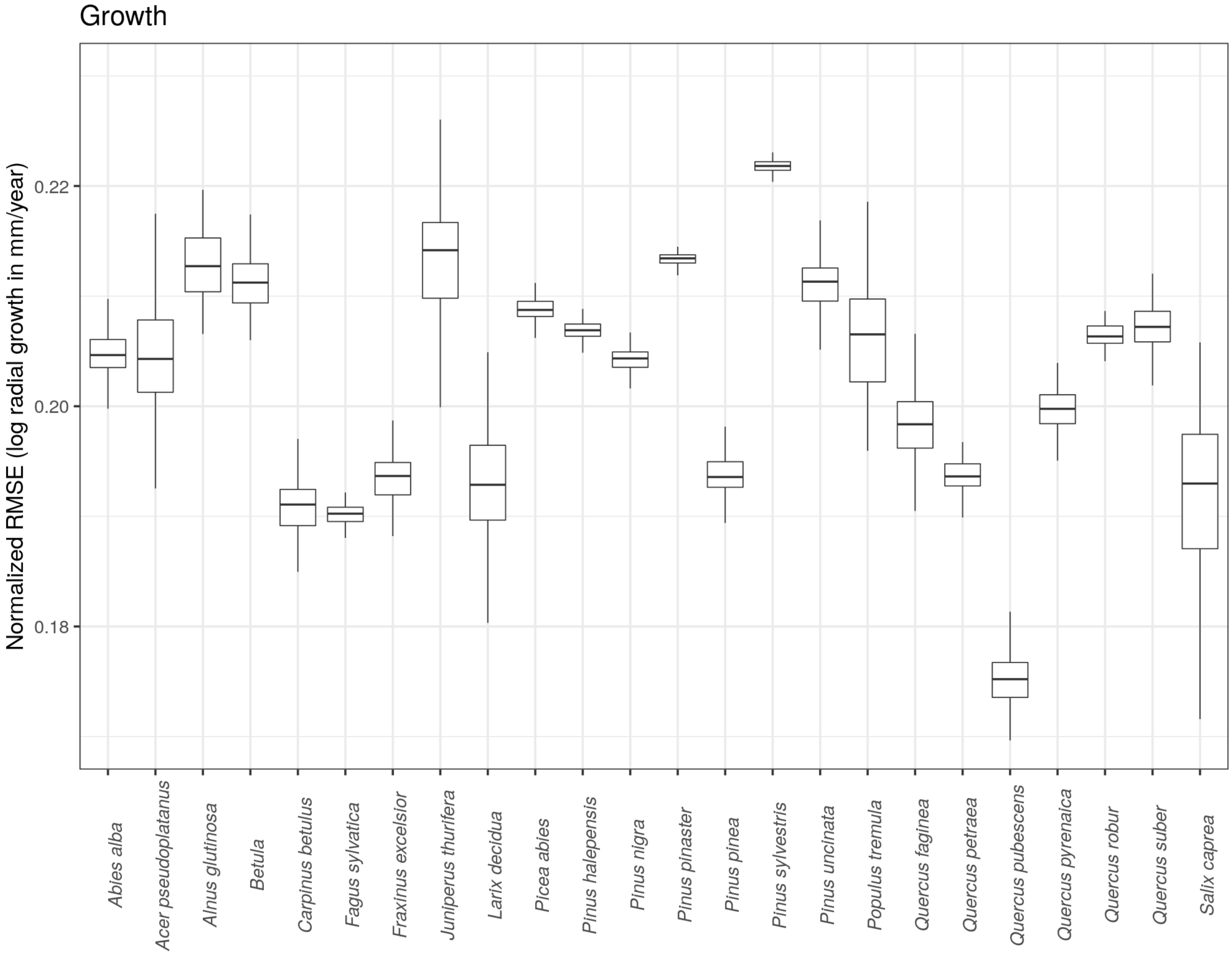

Figure S4: **Range of normalised root-mean-square error** over the 100 resampled growth models per species. Lower values of NRMSE indicate a better accuracy of the prediction on independent data.

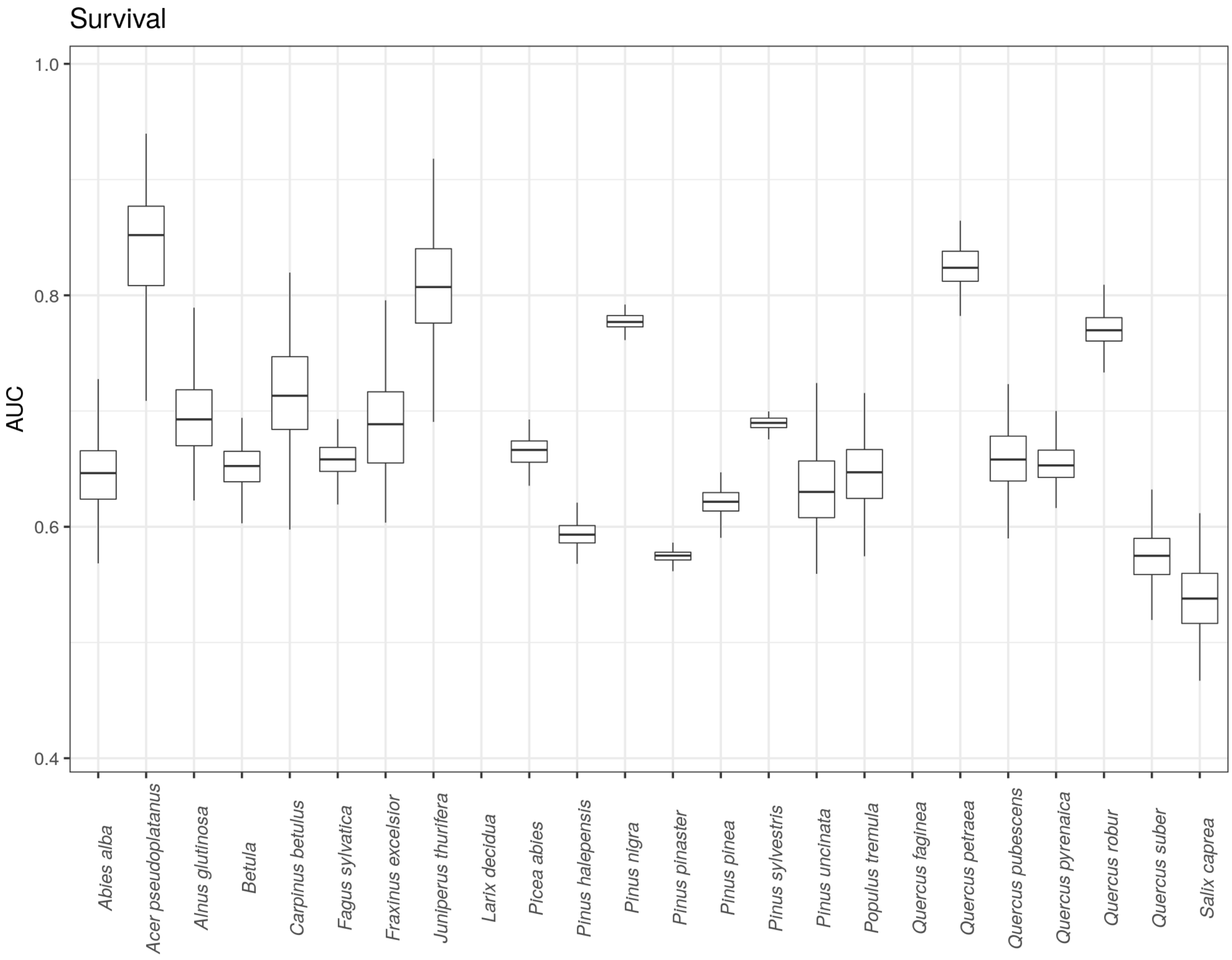

Figure S5: **Range of AUC over the 100 resampled survival models per species**. A higher AUC indicates better discriminatory capacity of the survival model on independent data.

#### 2.3 Recruitment function

**Aim** Our objective was to model the number of new recruits as a function of climate (using similar formulations as for growth and survival, without interactions), the abundance of adult conspecific, and competition through stand basal area. We used heterospecific basal area and conspecific basal area to represent two different effects of surrounding trees on recruitment: conspecific basal area will likely increase the expected number of recruits via seed production, while both conspecific and heterospecific basal area will decrease the number of recruits through competitive effects.

**Recruitment data** In our NFI datasets, we used ingrowth trees (trees that grew larger than the 10 cm dbh threshold between two surveys) as recruitment observations. To fit the recruitment functions, we excluded individuals recruited with a dbh $\geq$ 20 cm. We considered that new trees with this size corresponded to protocol artefacts: *e.g.* trees that were not measured in the first survey because they were located in a larger subplot where the minimum dbh was above 10 cm (see details on the protocol of each country in Table S1).

##### 2.3.1 Selection of the distribution

Count data are generally modelled with Poisson distributions. In our case the zero-inflation of the data (more zeroes than expected by a Poisson) led to poor fit, we thus considered other distributions. Zero-inflated Poisson models may be used to correct this bias of too many zeroes, it however leads to an increase in the number of parameters without strong process-based explanation for the zero-inflated part of the model. We also considered negative binomial distributions, which add a degree of liberty (overdispersion parameter) when compared to the Poisson model. Its main advantage is to keep the number of parameters relatively low, but still allow a better match with the observed distribution (especially the number of zeroes). Zell, Rohner, Thürig, and Stadelmann (2019) concluded that zero-inflated Poisson models gave only slightly better results than negative binomial (R2 of 30.9 against 30.00) when fitted to recruitment from the Swiss NFI data, but needed 22 parameters against 12. Zhu, Woodall, Monteiro, and Clark (2015) also used zero-inflated Poisson, without testing different distributions.

We conducted several tests to select the distribution, while also testing the model formulation (see next paragraph). Using a model based on equation S6 (see below), we evaluated the goodness of fit of the model for three distributions (Negative binomial, Poisson and zero-inflated Poisson) with three metrics:

1. the AIC (Akaike information criterion) of the model fitted using the whole dataset
2. the out of sample log likelihood (mean log likelihood calculated for 10 resampled model trained on 70% of the dataset on the remaining 30%)
3. the R2 calculated using the predicted response ($R2 = 1 - \frac{residuals}{variance(observation)}$ using the whole dataset)

Results (see Table S3) show slightly lower R2 when choosing the negative binomial distribution (here R2 only accounts for the mean prediction), but systematic higher out of sample log likelihood and lower AIC. We also compared the simulated distributions with the fitted models (by drawing the number of simulated recruits for each plots from the fitted distribution) to the observed distributions. More specifically, we compared the simulated *vs.* observed number of plots with n recruits (n ranging from zero recruit, one recruit, two recruits, up to the maximum number of recruits). We summarised these results for all species in Figure S6, with the correlation coefficient and the slope of simulated *vs.* observed frequencies distribution of the number of recruits per plots. The correlation and slope are equal to one when distributions are identical.

Results clearly show that the agreement between distributions of predicted recruitment count and observed recruitment count is better for the negative binomial model.

| **Species** | **AIC on whole dataset** | | | **Log likelihood on test dataset** | | | **R2** | | |
|---|---|---|---|---|---|---|---|---|---|
| | NB | Poisson | ZI Poisson | NB | Poisson | ZI Poisson | NB | Poisson | ZI Poisson |
| *Abies alba* | 5611 | 6333 | 5860 | -859 | -983 | -906 | 0.19 | 0.21 | 0.22 |
| *Acer campestre* | 5809 | 6410 | 6068 | -867 | -957 | -907 | 0.19 | 0.20 | 0.20 |
| *Acer pseudoplatanus* | 3424 | 3838 | 3517 | -508 | -571 | -525 | 0.14 | 0.14 | 0.15 |
| *Alnus glutinosa* | 4666 | 5974 | 5343 | -694 | -888 | -797 | 0.12 | 0.12 | 0.12 |
| *Betula* | 28460 | 37136 | 32837 | -4285 | -5635 | -4967 | 0.10 | 0.10 | 0.10 |
| *Capinus betulus* | 12327 | 13539 | 12652 | -1836 | -2014 | -1887 | 0.07 | 0.07 | 0.07 |
| *Fagus sylvatica* | 18277 | 21481 | 19546 | -2748 | -3225 | -2940 | 0.22 | 0.23 | 0.23 |
| *Fraxinus excelsior* | 10866 | 12263 | 11380 | -1623 | -1845 | -1709 | 0.10 | 0.11 | 0.11 |
| *Juniperus thurifera* | 4303 | 4865 | 4595 | -650 | -756 | -706 | 0.13 | 0.14 | 0.13 |
| *Larix decidua* | 835 | 969 | 870 | -129 | -151 | -137 | 0.09 | 0.12 | 0.12 |
| *Picea abies* | 26456 | 36573 | 31456 | -3968 | -5587 | -4783 | 0.13 | 0.14 | 0.14 |
| *Pinus halepensis* | 27605 | 34246 | 30717 | -4164 | -5177 | -4647 | 0.08 | 0.09 | 0.09 |
| *Pinus nigra* | 25372 | 38015 | 32345 | -3827 | -5760 | -4898 | 0.14 | 0.14 | 0.14 |
| *Pinus pinaster* | 27737 | 44680 | 35603 | -4147 | -6640 | -5265 | 0.08 | 0.09 | 0.08 |
| *Pinus pinea* | 7325 | 10253 | 8424 | -1102 | -1549 | -1272 | 0.05 | 0.05 | 0.05 |
| *Pinus sylvestris* | 60068 | 88712 | 76614 | -9038 | -13449 | -11599 | 0.16 | 0.16 | 0.16 |
| *Pinus uncinata* | 3588 | 4811 | 4377 | -537 | -732 | -669 | 0.24 | 0.27 | 0.26 |
| *Populus tremula* | 6468 | 7709 | 6994 | -975 | -1170 | -1062 | 0.14 | 0.15 | 0.15 |
| *Prunus padus* | 4784 | 5337 | 4992 | -722 | -801 | -754 | 0.07 | 0.07 | 0.09 |
| *Quercus faginea* | 11063 | 13943 | 12570 | -1662 | -2120 | -1906 | 0.08 | 0.08 | 0.09 |
| *Quercus ilex* | 40192 | 49293 | 42436 | -6084 | -7560 | -6416 | 0.20 | 0.26 | 0.28 |
| *Quercus petraea* | 15316 | 17409 | 15993 | -2298 | -2629 | -2403 | 0.21 | 0.22 | 0.23 |
| *Quercus pubescens* | 13773 | 15136 | 14453 | -2070 | -2299 | -2184 | 0.36 | 0.37 | 0.37 |
| *Quercus pyrenaica* | 13006 | 19516 | 16228 | -1955 | -3002 | -2477 | 0.12 | 0.13 | 0.13 |
| *Quercus robur* | 23946 | 27824 | 25267 | -3612 | -4196 | -3801 | 0.16 | 0.16 | 0.16 |
| *Quercus suber* | 8066 | 10648 | 8990 | -1199 | -1601 | -1340 | 0.11 | 0.16 | 0.16 |
| *Salix caprea* | 5334 | 6294 | 5797 | -804 | -956 | -887 | 0.08 | 0.08 | 0.08 |

Table S3: **AIC, Out of sample log likelihood and mean prediction R2 on the whole dataset for 3 different distributions** (NB: Negative binomial, Poisson and ZI Poisson:zero-inflated Poisson)

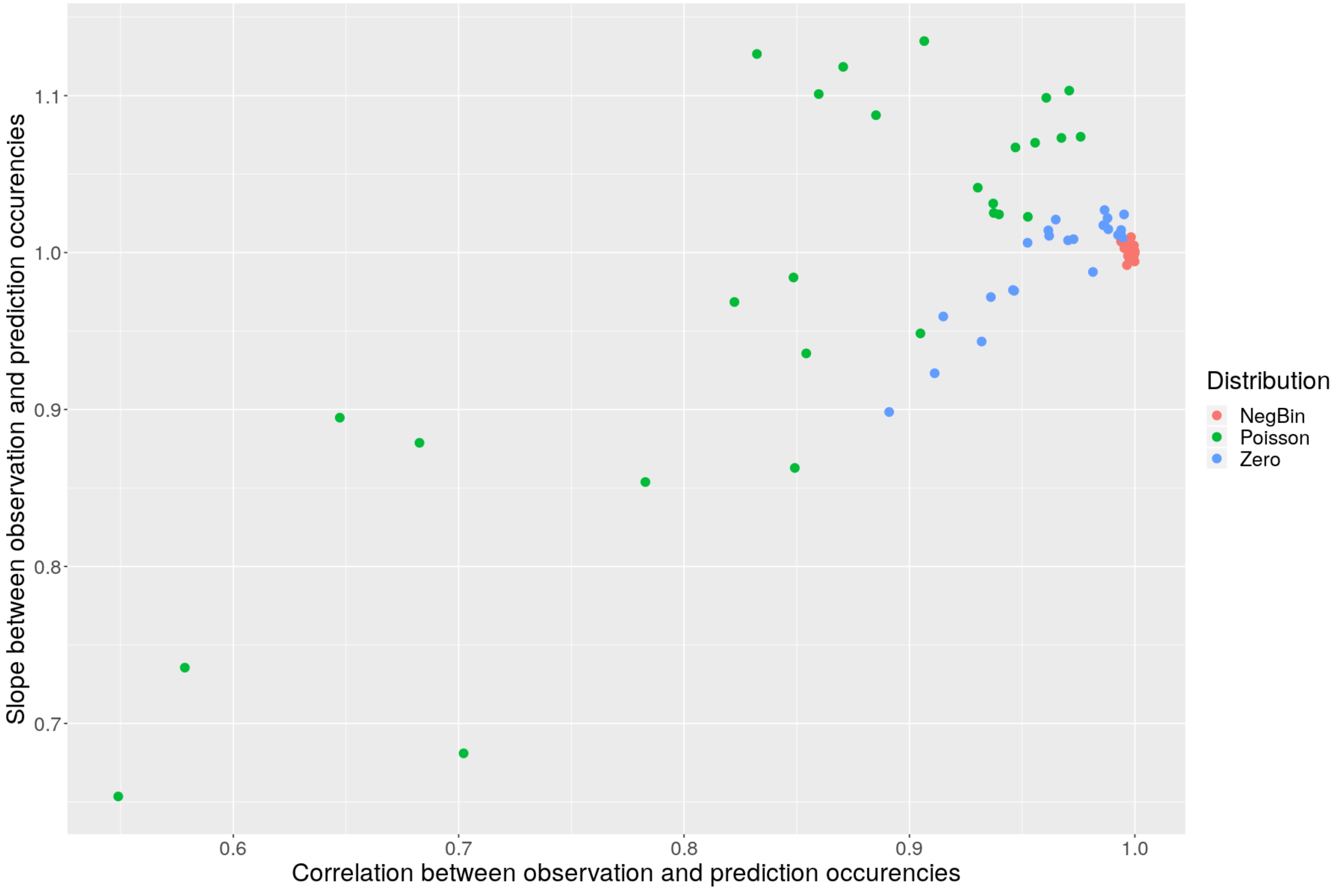

Figure S6: **Comparison of the simulated and the observed number of recruits per plot with three different type of distributions used to model recruit count: Negative binomial (red), Poisson (green) and zero-inflated Poisson (blue) distribution**. We evaluated goodness of fit for each species by comparing the simulated and the observed number of plots with n recruits using two metrics, the correlation (x-axis) and slope (y-axis) between observed and simulated number of plots with n recruits for each species. Each point represents one type of distribution for one species.

##### 2.3.2 Selection of model shape and covariates

As for growth and survival, different formulations of climate response curves were tested (polynomial as in equation S3 or asymptotic as in equation S2, represented for simplicity as $f(clim_i)$ in the equation S6).

Details on the processes of seed production, germination and seedlings growth/survival were not included in the formulation because they were not observed in the NFI data. To represent the positive effect of the abundance of conspecifics on seed availability and the negative effect of intra- and interspecific competition, the local stand structure is summarised with two variables: the basal area of conspecific individuals ($BA^{con}$), and the basal area of heterospecific individuals ($BA^{con}$). The shape of recruitment rate versus basal area of both conspecific and heterospecific is relevant, as it will have an impact on the equilibrium properties of the system (see Eager, Rebarber, & Tenhumberg, 2012; Townley, Rebarber, & Tenhumberg, 2012, for the specific case of Integral Projection Model). For numerous plant species, density dependence is expressed as $f(x) = g(x) * x$, x being a proxy for species abundance, here the basal area, $g$ being generally decreasing and non-negative. This formulation is classic in ecology, similar to the model of Lines, Zavala, Ruiz-Benito, and Coomes (2020) for juvenile recruitment. We tested several formulations to model recruitment count, each being based on a count data distribution, with a log link. For each species, we fitted six alternative models of the density dependence effect of basal area (see Table S4).

We repeated these fits 100 times using a sample of 70% of the available dataset and computed the AIC on the remaining 30%. We then selected the model that resulted in the best mean AIC across all species and climatic

| Model | Equation |
|---|---|
| 1 | $log(Rec_p) = K_c + \alpha * log(BA_p^{con}) + \beta * BA_p^{con} + f(clim_p) + log(time_p)$ |
| 2 | $log(Rec_p) = K_c + \alpha * log(BA_p^{con}) + \beta * BA_p^{tot} + \gamma * BA_p^{het} + f(clim_p) + log(time_p)$ |
| 3 | $log(Rec_p) = K_c + \alpha * log(BA_p^{con}) + \beta * BA_p^{con} + \gamma * BA_p^{het} + f(clim_p) + log(time_p)$ |
| 4 | $log(Rec_p) = K_c + \beta * BA_p^{con} + \gamma * BA_p^{het} + f(clim_p) + log(time_p)$ |
| 5 | $log(Rec_p) = K_c + \alpha * log(BA_p^{con}) + \beta * BA_p^{con} + \gamma * BA_p^{het} + \delta * BA_p^{con} * BA_p^{het} + f(clim_p) + log(time_p)$ |
| 6 | $log(Rec_p) = K_c + \alpha * log(BA_p^{con}) + \beta * (BA_p^{con})^2 + \beta * BA_p^{con} + \gamma * BA_p^{het} + f(clim_p) + log(time_p)$ |

Table S4: **Alternative models explored for the density dependence effect.**

dependencies. Finally, we also used GAMs (Generalized Additive Models) to verify that the shape of the density-dependence function of our parametric model matched well the data. The structure of the selected recruitment model is:

$$log(Rec_p) = K_c + \alpha * log(BA_p^{con}) + \beta * BA_p^{con} + \gamma * BA_p^{het} + f(clim_p) + log(time_p) \quad \text{(S6)}$$

where $Rec_p$ is the observed number of recruits in plot $p$, $K_c$ is mean the number of recruits per country (to account for differences in protocols between countries), $\alpha$, $\beta$, and $\gamma$ are estimated parameters and $f(clim_p)$ the function describing the climatic response to climate in plot $p$, $sgdd_p$ and $wai_p$, with the shapes described above for the growth and survival model. $log(time_p)$ is included as an offset to account for the effect of the time between census on the observed number of recruits. For the IPM simulations we used the average of the intercept across countries.

It is worth noticing that this formulation excludes recruitment in plots where a species is initially absent; these plots have therefore been excluded from the analysis (this represents about 9% of the data).

##### 2.3.3 Recruitment response curves

Figure S7 represents the recruitment at the different tree niche positions (hot edge, center, and cold edge) as a function of conspecific basal area. Heterospecific basal area was set to zero in this figure. The parameters of the 100 refitted models of each species are provided in '.rds' R object in the Zenodo repository https://zenodo.org/record/6424490 (for instance for *Fagus sylvatica* in the file 'output/fit_sgr_all_Fagus sylvatica.Rds').

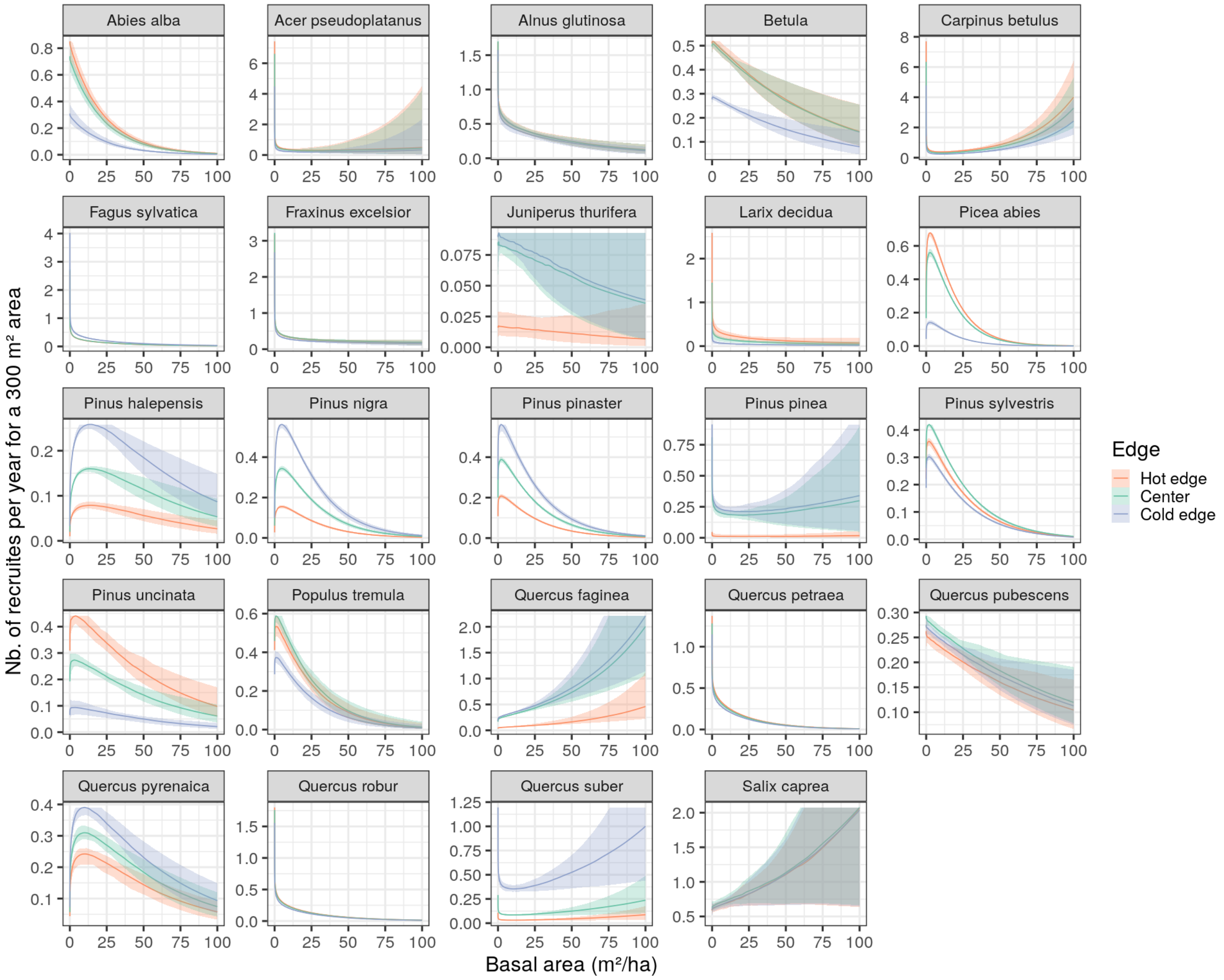

Figure S7: **Recruitment rates according to stand basal area** (conspecific). Each color corresponds to a climatic niche position, lines represent the median value between all 100 sub-models, and the shaded areas the interval between the 5th percentile and the 95th percentile.

##### 2.3.4 Recruitment goodness of fit

We cross-validated the 100 recruitment models fitted on a re-sampling of 70% of the data on the remaining 30% by computing the normalised root-mean-square error (NRMSE) (Figure S8). We also validated that the number of plots predicted with zero recruits matched well the observed number of plots with zero recruits per species (Figure S9).

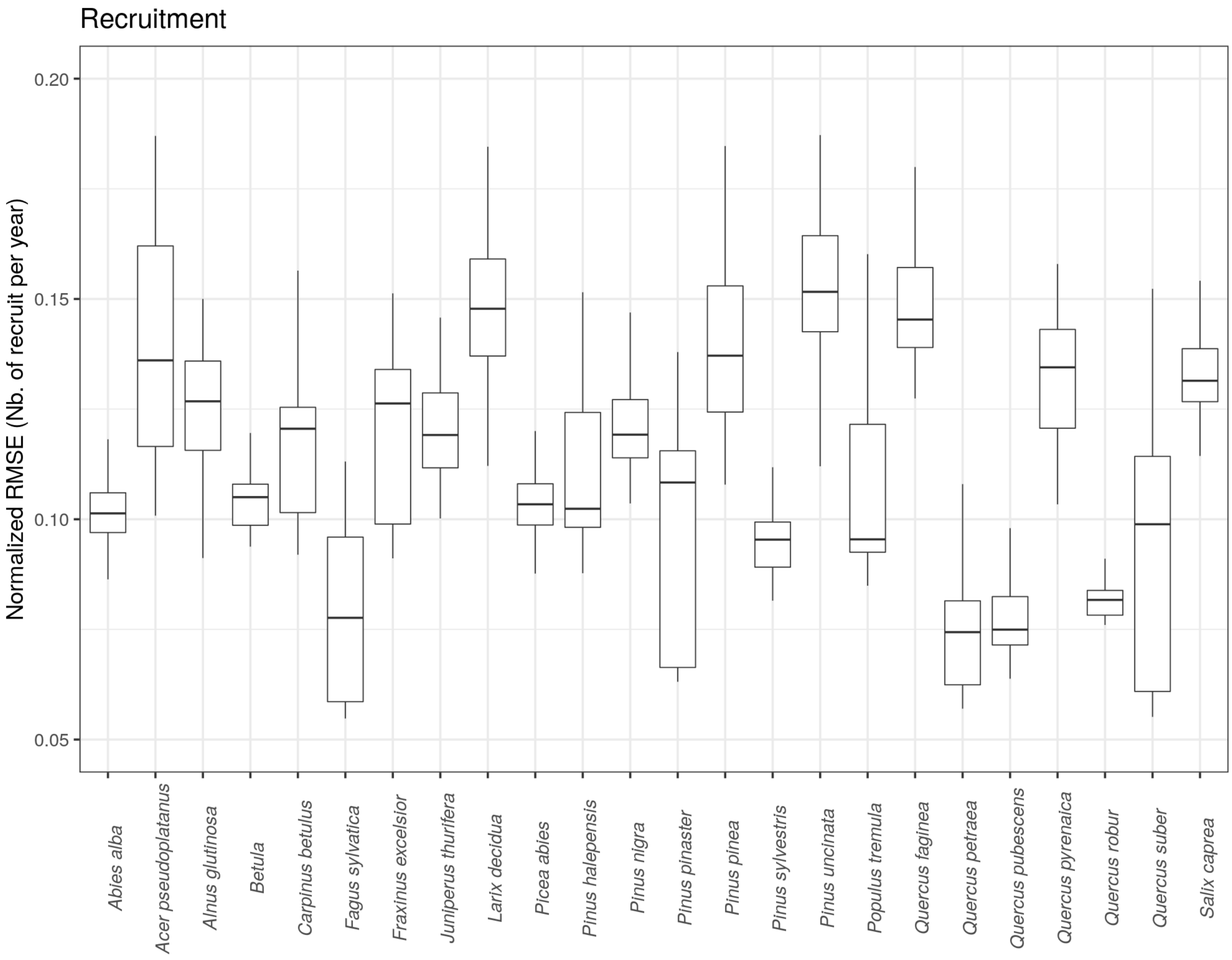

Figure S8: **Range of normalised root-mean-square error** over the 100 resampled recruitment models per species. Lower values of NRMSE indicate a better accuracy of the prediction on independent data.

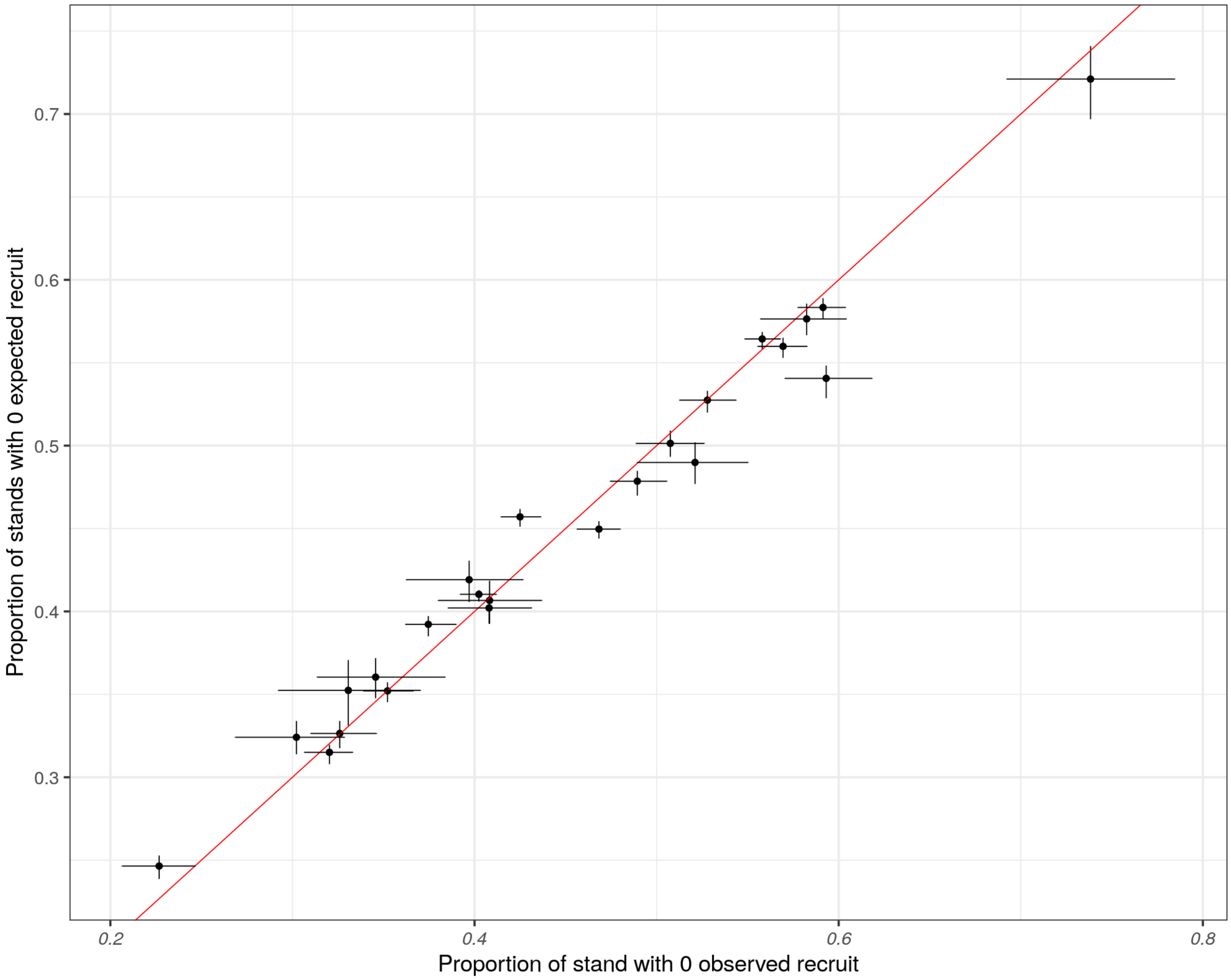

Figure S9: **Predicted proportion of plots with zero recruits vs observed proportion of plots with zero recruits per species.**

#### 2.4 IPM matrix formulation

Equation (1) can be re-written in a matrix format as :

$$\mathbf{X_{t+1}} = (\mathbf{P} + \mathbf{F}) \times \mathbf{X_t} \tag{S7}$$

with $\mathbf{X_t}$ and $\mathbf{X_{t+1}}$ being the discretized size distribution (number of individuals per size classes, corresponding to integration of $n(z,t)$ over each size class) at time $t$ and $t+1$, respectively; $\mathbf{P}$ is the transition matrix accounting for the growth and survival, corresponding to the kernel $P(z',z)$ and $\mathbf{F}$ is the fecundity matrix corresponding to the kernel $F(z',z)$.

Due to the density dependence of growth, survival, and recruitment rates, equation S7 is actually written as

$$\mathbf{X_{t+1}} = \left(\mathbf{P}(BA_t^{tot}) + \mathbf{F}(BA_t^{het}, BA_t^{con})\right) \times \mathbf{X_t} \tag{S8}$$

$BA_t^{tot}$ is the total basal area at time t and is divided in two components in the fecundity matrix: heterospecific and conspecific basal area such that $BA_t^{tot} = BA_t^{het} + BA_t^{con}$.

We included a delay in tree recruitment to account for the time it takes for a sapling to reach the minimum dbh, meaning that a newly recruited tree is integrated into the population only after 10 years. We chose to use a constant value of 10 years across species and climatic conditions to simplify the model because we had no direct data on this parameter. 10 years is a relatively short time for trees to reach the minimum dbh, corresponding to the faster growing species, but this allows to reduce simulation time. Preliminary exploration of changes in the length of this delay showed that this had little impact on the response of invasion rate at the edge. Indeed, we compare climatic edges and climatic center with the same delay, which probably limits the importance of this parameter.

##### 2.4.1 Building the fecundity matrix

Following Townley et al. (2012), we built the fecundity matrix $\mathbf{F}$ from the recruitment function $f_{rec}$ relating the number of recruits to conspecific and heterospecific basal area:

$$\begin{aligned}
\mathbf{X}_{t+1}^{recruits} &= f_{rec}(BA_t^{Het}, BA_t^{con})\mathbf{b} \\
&= \frac{f_{rec}(BA_t^{Het}, BA_t^{con})}{BA_t^{con}}\mathbf{b}.(\mathbf{c}.\mathbf{X_t}) \\
&= \mathbf{F}(BA_t^{Het}, BA_t^{con}).\mathbf{X_t}
\end{aligned} \tag{S9}$$

with $\mathbf{F}(BA_t^{Het}, BA_t^{con}) = \dfrac{f_{rec}(BA_t^{Het}, BA_t^{con})}{BA_t^{con}}\mathbf{b}.\mathbf{c}$

where $\mathbf{b}$ is the vector representing the size distribution of recruits, and $\mathbf{c}$ is a vector of weights built such that $\mathbf{c}\mathbf{X_t} = BA_t^{con}$.

##### 2.4.2 Numerical integration of the kernel

To compute the transition matrix $\mathbf{P}$, we integrated the probability distribution of the growth kernel $P$. We used a Gauss-Legendre integration method to perform integration close to the diagonal (corresponding to a radial growth below 5 $cm$ per year) where the bulk of the kernel is concentrated, and a mid-bin integration for the rest of the IPM where the growth kernel is less variable and close to 0. We used the 'gauss.quad' function from the R package 'statmod' with 140 nodes for the integration on size at t+1, and 3 nodes for the integration of size at time t (420 per bins for integration on the two dimensions).

For each species, we divided the dbh range into 700 bins, and we checked that integration was performed correctly by verifying that for each size class, the distribution of size at time step $t+1$ integrates to the survival probability of this size class (Easterling, Ellner, & Dixon, 2000). We also verified that the invasion rate metric (calculated at low competition and at the climatic niche center) is close to its asymptotic value (see Figure S10) when choosing 700 bins, with a maximal error below 1 % and a median error below 0.1%.

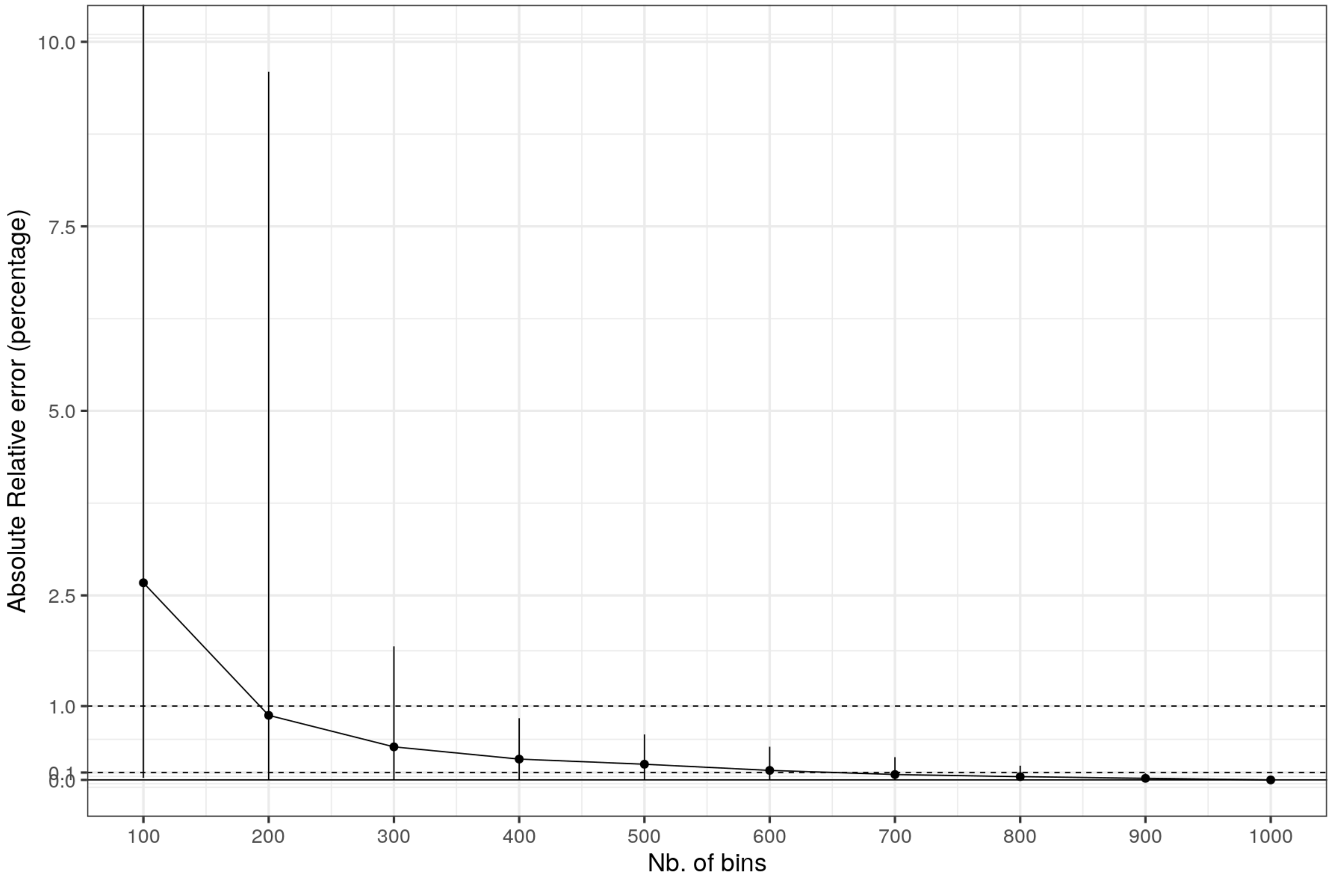

Figure S10: **Variation of the absolute relative error in the invasion rate at low competition using different numbers of bins for integration of the IPM**. For each number of bins, the invasion rate is computed for all species x model combinations. Points represent the median of the relative error across species x model combination, and vertical lines the range between relative errors.

IPM dynamics may lead to eviction, i.e. an individual may grow above the maximum of the IPM size. This concerns the upper size classes, where growth will lead to individuals reaching a size above the maximum size of the IPM. We need to consider a correction for growth integration in this case; we chose to suppress all trees reaching the last class size, thus assuming that trees die above this size (maximum size for each species is 1.1 times the maximum observed dbh).

#### 2.5 Accounting for eviction

Williams, Miller, and Ellner (2012) showed that there is no universal solution to solve eviction, but rather the best solution has to be selected based on the source of eviction in the IPM and the specificity of the biological system modeled with the IPM. In our case, eviction occurs because (1) the unlimited growth and (2) the tree longevity (Munné-Bosch, 2018; Piovesan & Biondi, 2021). First, for some species tree radial growth declines at large size but never reaches zero, leading to a steady increase in tree size over time. Second, although our statistical model captures a decline in survival with size for most species (see Figure S2), we likely underestimate this decline. Reasons are (1) the rarity of very large individuals in the observations and (2) the removal of senescent trees by management (Munné-Bosch, 2018). In Kunstler et al. (2021) we showed that from a cohort starting at the minimum dbh, only a small proportion of large trees continue to grow and survive above the maximum observed diameter for the species. In Kunstler et al. (2021) we thus decided to use the ceiling eviction correction proposed by Williams et al. (2012) which essentially consists of adding an additional size class to keep track of the few individuals that grow over the maximum size during the cohort lifetime. In this new analysis, this approach was not feasible because we run the simulation over a much longer time frame and the small percentage of individuals growing over the maximum size class accumulates over the generations. We thus decided to rather implement a different correction where we assumed that all trees that grow over the maximum size class (defined as 1.1 of the maximum observed dbh) are killed. This assumption is implemented in numerous models of forest dynamics, usually via a species-specific maximum age (see Bugmann et al. (2019)).

#### 2.6 Equilibrium state

For the case where only the recruitment rate is density-dependent (the transition matrix $\mathbf{P}$ is not density-dependent), Rebarber, Tenhumberg, and Townley (2012) and Townley et al. (2012) proposed analytical solutions to compute equilibrium stability conditions. In this case, the IPM is defined as:

$$\mathbf{X_{t+1}} = \mathbf{P}\mathbf{X_t} + \mathbf{b}f(\mathbf{c}\mathbf{X_t}) \tag{S10}$$

where matrix $\mathbf{P}$ is non-negative and has a spectral radius strictly below 1, $\mathbf{b}$ and $\mathbf{c}$ are positive vectors and f is a real function.

The equilibrium state for this IPM model, with $\mathbf{P}$ constant, is defined by the population vector $\mathbf{X_e^*}$:

$$\mathbf{X_e^*} = (\mathbf{Id} - \mathbf{P})^{-1}.p_e^*.BA_e^* \tag{S11}$$

where $p_e^* = \dfrac{1}{\mathbf{c}.(\mathbf{Id}-\mathbf{P})^{-1}.\mathbf{b}}$, $BA_e^*$ is the solution of $f_{rec}(BA_e^*) = p_e^*.BA_e^*$, and $\mathbf{Id}$ is the identity matrix. We use the notation $\mathbf{X_e^*}$ and $BA_e^*$, to make the distinction between the analytical solution of equilibrium in the case that $\mathbf{P}$ is constant and the equilibrium $\mathbf{X_e}$ and $BA_e$ derived with simulations accounting for the density dependence in $\mathbf{P}$.

Based on equation S11, if we compute a fixed transition matrix $\mathbf{P}$ for constant basal area (noted $BA_{cst}$) we can then derive the equilibrium vector $X_e^*$ and $BA_e^*$. Using this approach, we verified that the value of $\mathbf{X_e}$ and $BA_e$ identified by simulations as described in the main text lead to $BA_e^* = BA_e$ when $\mathbf{P}$ is computed at a constant basal area value $BA_{cst} = BA_e$. This gives an analytical confirmation that an equilibrium identified by simulations is a true equilibrium.

As described in the main text, we excluded simulations when equilibrium tree density was either below 1 $m^2ha^{-1}$ or above 200 $m^2ha^{-1}$. The Table S5 gives the number of simulations excluded and $BA_e$ per species x edge combination. Out of 8100 simulations (3 climatic conditions * 27 species * 100 resampling), 68 went below 1 $m^2ha^{-1}$, and 647 reached BA values above 200 $m^2ha^{-1}$. Overall, 717 simulations were discarded (about 9%). We excluded species in climatic conditions if less than 50% of simulations reached equilibrium, which lead to the exclusion of *Acer campestre*, *Pinus pinea* and *Prunus padus* at the hot edge, and *Quercus faginea* and *Quercus ilex* at the cold edge. *Pinus pinea* at the hot edge was the only exclusion due to too low BA values at equilibrium.

#### 2.7 Individual-based simulations

We performed IBM simulations by first fixing the sampling area to 1 $ha$ to compute demographic stochasticity, or to 0.1 $ha$ to compute extinction time. We drew the number of individual trees for each bin size from the distribution previously computed at equilibrium $X_e$, and attributed to each tree a dbh equal to the mean dbh of its bin. We directly used the vital rate functions to draw a growth value, individual survival event, and the number of recruits at time $t+1$ from the IPM kernels.

#### 2.8 Metrics calculation

**Invasion rate** To evaluate mean population performance, we used the invasion rate. In size-structured populations, the invasion rate is measured by the net reproductive rate, $R_0$, of a rare invader (Falster, Brännström, Westoby, & Dieckmann, 2017). For density independent cases, $R_0$ is estimated by the dominant eigenvalue of the matrix $\mathbf{F}.(\mathbf{I} - \mathbf{P}^{-1})$ (Ellner et al., 2016).

We approximated $R_0$ in our density-dependent IPM by simplifying the recruitment function of the invading species when rare as a constant fecundity matrix. We considered $F$ fixed, corresponding to a constant level of heterospecific competition $BA_t^{het}$, and conspecific competition ($BA_t^{con}$). To compute the net reproductive rate when the conspecifics were rare, we set the conspecific basal area to a low value of 0.1 $m^2ha^{-1}$ in $F$. We did not account for the competitive effect of conspecific basal area on fecundity. Furthermore, we fixed the transition matrix $\mathbf{P}$ at

| Species | Nb. of resampled models not reaching equilibrium | Mean basal area $m^2ha^{-1}$ | Tree density $Nbha^{-1}$ |
|---|---|---|---|
| | hot/center/cold | | |
| *Pinus sylvestris* | 0/0/0 | 28/31/32 | 374/408/390 |
| *Pinus pinaster* | 0/0/0 | 17/23/22 | 289/348/369 |
| *Picea abies* | 0/0/0 | 37/36/9 | 253/275/343 |
| *Pinus nigra* | 0/0/0 | 31/40/49 | 339/356/372 |
| *Pinus halepensis* | 0/0/0 | 4/23/44 | 167/397/560 |
| *Quercus ilex* | 0/0/90 | 10/49/NA | 143/1286/NA |
| *Fagus sylvatica* | 0/0/0 | 41/43/42 | 279/291/494 |
| *Quercus robur* | 0/0/0 | 29/33/39 | 241/233/229 |
| *Betula* | 0/0/0 | 35/36/16 | 829/870/537 |
| *Quercus petraea* | 0/0/0 | 32/32/34 | 216/235/266 |
| *Quercus pubescens* | 0/0/0 | 28/36/33 | 556/637/684 |
| *Quercus pyrenaica* | 0/0/0 | 46/42/44 | 593/657/899 |
| *Quercus suber* | 0/1/14 | 14/38/58 | 111/517/2312 |
| *Pinus pinea* | 65/0/0 | 0/17/17 | 0/278/277 |
| *Abies alba* | 0/0/0 | 47/49/33 | 285/277/254 |
| *Carpinus betulus* | 3/16/48 | 64/89/97 | 2689/5500/5768 |
| *Pinus uncinata* | 0/0/0 | 30/36/36 | 775/662/319 |
| *Fraxinus excelsior* | 0/0/0 | 47/52/45 | 588/599/541 |
| *Quercus faginea* | 21/24/52 | 82/98/125 | 2114/6027/10064 |
| *Alnus glutinosa* | 0/0/0 | 22/38/58 | 527/554/725 |
| *Juniperus thurifera* | 18/0/1 | 20/20/21 | 93/331/389 |
| *Populus tremula* | 0/0/0 | 34/35/23 | 403/408/390 |
| *Acer campestre* | 99/100/97 | NA/NA/NA | NA/NA/NA |
| *Acer pseudoplatanus* | 0/0/14 | 57/70/63 | 1112/1639/2366 |
| *Larix decidua* | 0/0/0 | 34/26/20 | 256/194/106 |
| *Salix caprea* | 1/0/0 | 28/30/39 | 1023/1196/1771 |
| *Prunus padus* | 51/0/0 | 72/50/33 | 1612/1117/625 |

Table S5: **Equilibrium conditions for the 27 analysed species**. For each species/climatic condition, 100 models were fitted with re-sampled data. In most cases, equilibrium was reached and basal area and tree density were computed. Models that did not reach equilibrium were discarded and the basal area and tree density were not derived (indicated by NA). We considered that a simulation didn't reach equilibrium if it leads to basal area less than $1m^2ha^1$ (as simulations work on continuous population abundances there is no strict extinction), or increasing above 200 $m^2ha^1$ (no upper limits to the population). Only trees above 10 cm of DBH are modelised. See the methods section in the main text for more details.

$BA_t^{tot} = BA_t^{het}$ as the abundance of the invading species is considered to be negligible when rare. We computed $R_0$ for two conditions of heterospecific competition: no heterospecific competition (where $BA^{het} = 0\ m^2ha^{-1}$), and a high level of heterospecific competition (where $BA^{het} = 60\ m^2ha^{-1}$).

**Time to extinction** Following Grimm and Wissel (2004), we estimated the parameter $T_m$, corresponding to the intrinsic mean time to extinction. This method is based on an estimation of the cumulative probability of population extinction at time $t$, $P_0(t)$ based on IBM simulations. We used 250 IBM simulations with a maximum of 3000 years for each resampled model, species and climatic condition to estimate $P_0(t)$. Then we plotted $-ln(1 - P_0(t)$ against $t$ and verified that it was in agreement with a linear regression, and extracted from the linear regression the intercept and the slope. $T_m$ is given by the inverse of the slope.

**Computation of demographic variance** To compute demographic variance, for each model we ran long-term individual-based stochastic simulations (3000 years). We set plot area to 1 $ha$, and initiated the number of trees for each size class from equilibrium tree density $X_e$. Following Engen, Lande, Sæther, and Dobson (2009) and Jaffré and Le Galliard (2016), we used the reproductive value $V$ to compute demographic variance as $Var(\Lambda_t|V_t)$ with $\Lambda_t = \dfrac{V_{t+1}}{V_t}$ and $V_t$ the reproductive value at time t. The reproductive value is computed using the population equilibrium structure derived from deterministic simulations: basal area is set constant at its equilibrium value, we then computed the scaled right eigenvector $\mathbf{v}$ of $\mathbf{P} + \mathbf{F}$, also called the reproductive value distribution, "which measures the contribution of an individual to future population growth relative to other individuals in the population" (Jaffré & Le Galliard, 2016). The total reproductive value is then computed as $V_t = \sum_{i=1}^{i=N} \mathbf{v}(i)\mathbf{X_t}(i)$.

**Damping time** Damping time, the time to converge to a stable size structure after a disturbance, is independent of the size structure of the perturbations. We computed the Jacobian matrix at $\mathbf{X_e}$ and extracted its eigenvalues. The real part of these eigenvalues were all negative, meaning the system was stable (Caswell, 2000). From the dominant eigenvalue $\lambda^1$, we derived a damping time computed as $Damp_{time} = \dfrac{log(2)}{Re(\lambda^1)}$.

**$T_0$ and $T_{half}$** We used simulations to derive the two metrics on short term recovery after a disturbance. For each species, we disturbed its population at demographic equilibrium (defined by its size distribution $X_e$ at equilibrium) by reducing the density of the largest trees (above the 66th percentile of dbh distribution at equilibrium) by a factor of two. This resulted in a decrease of tree density by $N_{pert}$. From these simulations, we extracted the time for the population to first return to its equilibrium density (regardless of the tree size distribution) - T0, and the time after which at least half of the initial tree reduction has been permanently recovered - Thalf ($T_{half}$ (as $N(t) > N_e - \frac{N_{pert}}{2}$).

The equations for all metrics are given in Table S6.

| Name | Computation |
|---|---|
| Invasion Rate<br>No competition | maximum eigenvalue of $\mathbf{F}(\mathbf{1}-\mathbf{P})^{-\mathbf{1}}$<br>with $\mathbf{F}$ and $\mathbf{P}$ computed at $BA^{het}=0$ |
| Invasion Rate<br>with competition | maximum eigenvalue of $\mathbf{F}(\mathbf{1}-\mathbf{P})^{-\mathbf{1}}$<br>with $\mathbf{F}$ and $\mathbf{P}$ computed at $BA^{het}=60$ |
| Nb. of individuals | $N_e=\sum_{i=1}^{i=700} Xe_i$<br>with **Xe** the size distribution at equilibrium |
| Demographic<br>Variance | $Var(\frac{V_{t+1}}{V_t}\|V_t)$<br>with $V_t$ the reproductive value at time t |
| Extinction Time $T_m$ | computed from regression -ln(1-$P_0(t)$) $\sim$ a + $\frac{t}{Tm}$<br>with $P_0(t)$ the cumulative probability of population extinction at time $t$ |
| Damping Time | $\frac{log(2)}{\lambda_1}$<br>with $\lambda_1$ the real part of the dominant eigen value<br>of Jacobian matrix computed at equilibrium |
| $T_0$ | time T at which $N(T)\approx N_e$<br>with $N(t)$ the tree density at time t,<br>and $N_e$ the tree density at equilibrium |
| $T_{half}$ | time T such as $\forall t>T$, $N(t)>N_e-\frac{N_{pert}}{2}$<br>with $N(t)$ the tree density at time t,<br>$N_e$ the tree density at equilibrium,<br>and $N_{pert}$ the initial perturbation |

Table S6: **Summary of the demographic metrics computed**.

### 3 Climatic edges and center

#### 3.1 Climatic axis

As described in the main text, to describe the species climatic distributions, we focused on the first axis of the PCA of the two climatic variables we used (see Figure S11). The two climatic variables are: (1) the sum of growing degree days above 5.5 °C (*sgdd*) and (2) water aridity index (*wai*) (their variation across the NFI plots are presented in Figure S12 and S13). The climatic data used in the analysis are provided in the Dryad Digital Repository https://doi.org/10.5061/dryad.wm37pvmkw (Ratcliffe et al., 2020).

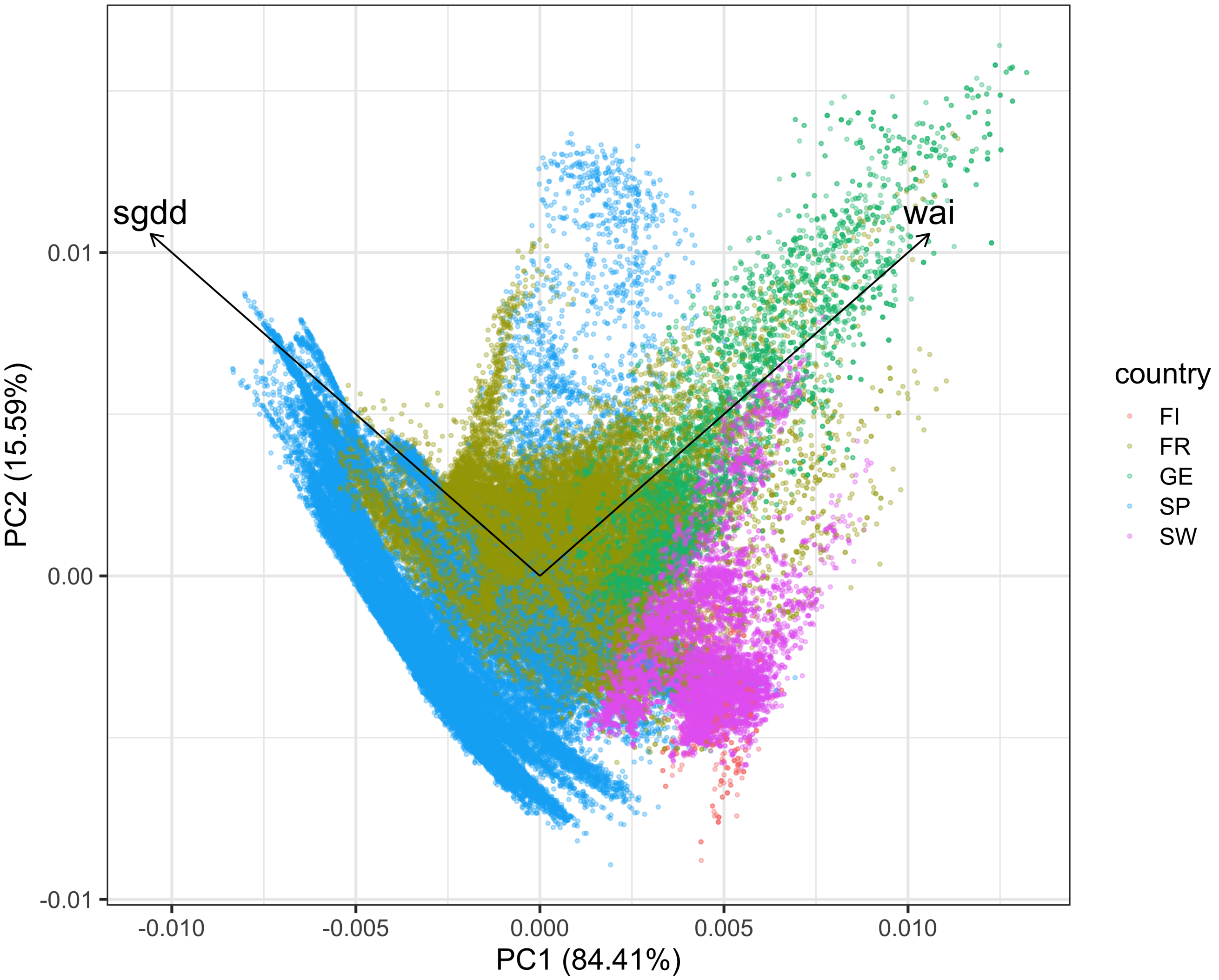

Figure S11: **PCA of the two selected climatic variables**, *sgdd* and *wai*. The percentage of variance explained by each axis is shown in the axis label. The country codes are: FI - Finland, FR - France, GE - Germany, SP - Spain, and SW - Sweden.

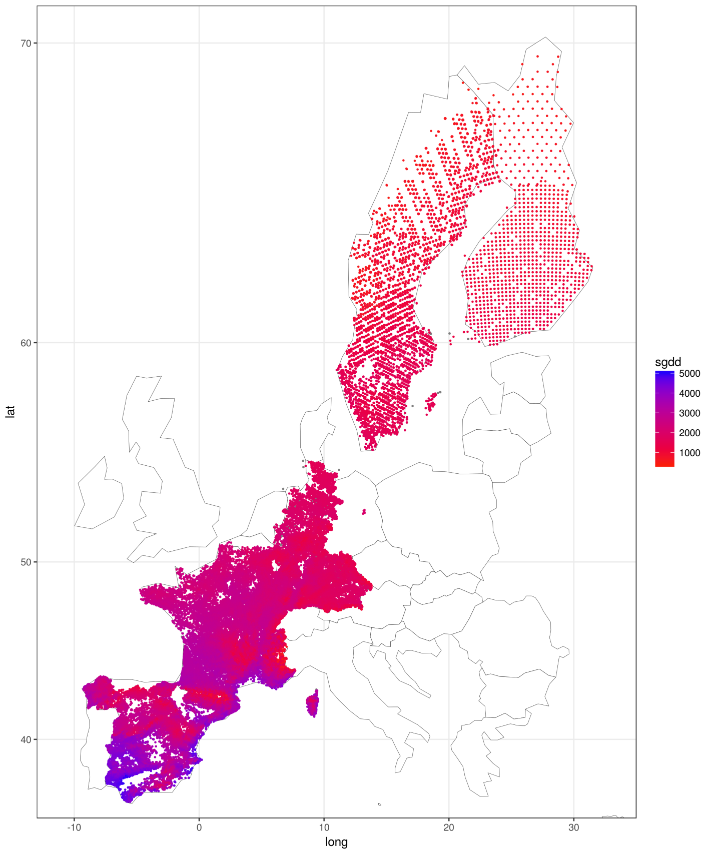

Figure S12: **Variation in the sum of growing days (sgdd) across FUNDIV NFI plots.**

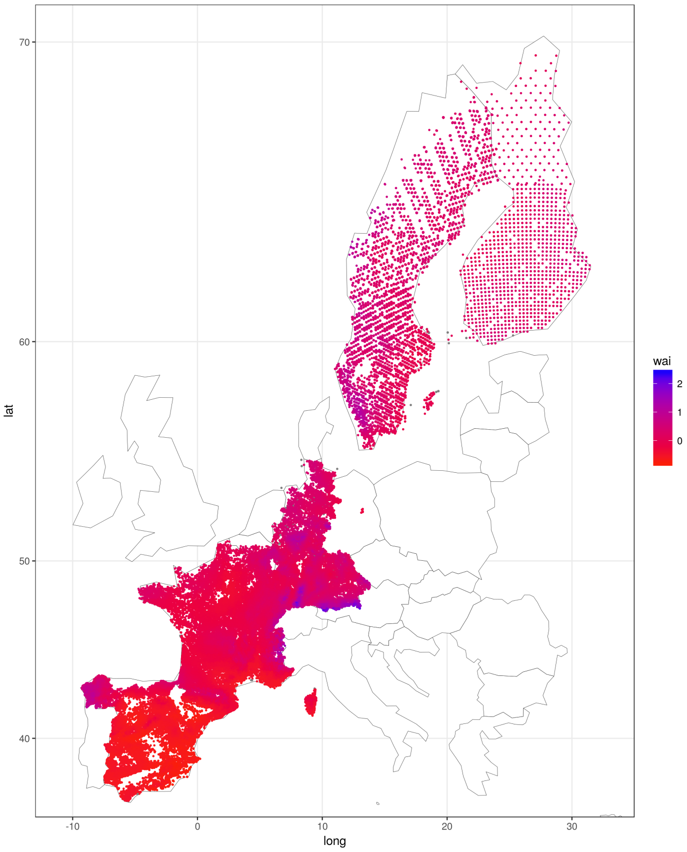

Figure S13: **Variation in the water aridity index (wai) across FUNDIV NFI plots.**

#### 3.2 Species distribution models

In order to evaluate which species edges in the FunDivEUROPE data correspond to an actual limit of the species distribution and not just to a limit in the coverage of the data, we fitted species distribution models with presence/absence data covering all Europe (Mauri, Strona, & San-Miguel-Ayanz, 2017) and retained only the edges for which there was a clear drop in the probability of presence of the species at the edge. We fitted ensemble species distribution models using four different models in BIOMOD2 (Thuiller, Lafourcade, Engler, & Araújo, 2009), using the EU-Forest data set (Mauri et al., 2017). This allowed us to have a robust estimation of the probability of presence of each species in each NFI plot. Then, we estimated the mean probability of presence at the center and the two edges by computing the mean probability of presence of the plots that were in climatic bins (as described by the *wai-sgdd* combination) corresponding to the edges or the center. We retained only edges with at least a 10 % drop in the probability of presence at the edge for comparison of the demographic performance at the edge *vs.* the center of the distribution.

##### 3.2.1 Species distribution data

Mauri et al. (2017) provides a synthesis of available data on European tree species distribution on a 1 x 1 km grid. A large part of this dataset overlaps with the NFI data we used, but it also includes NFI data from numerous other countries (including Italy, Norway, Austria, Switzerland, and several eastern European countries) and other data sources, such as the forest focus data base and the Biosoil data base corresponding to a total of 1,000,525 occurrence records.

The FunDivEUROPE NFI dataset only provides a genus-level description for *Betula*, as species-level information are not available in all countries. We thus derived a mean prediction for the two *Betlua* species available in EU-Forest *Betula pendula* and *Betula pubescens*.

#### 3.3 Species distribution and edges selection

The figure S14 presents the distribution of the species and the edges selected for the analysis.

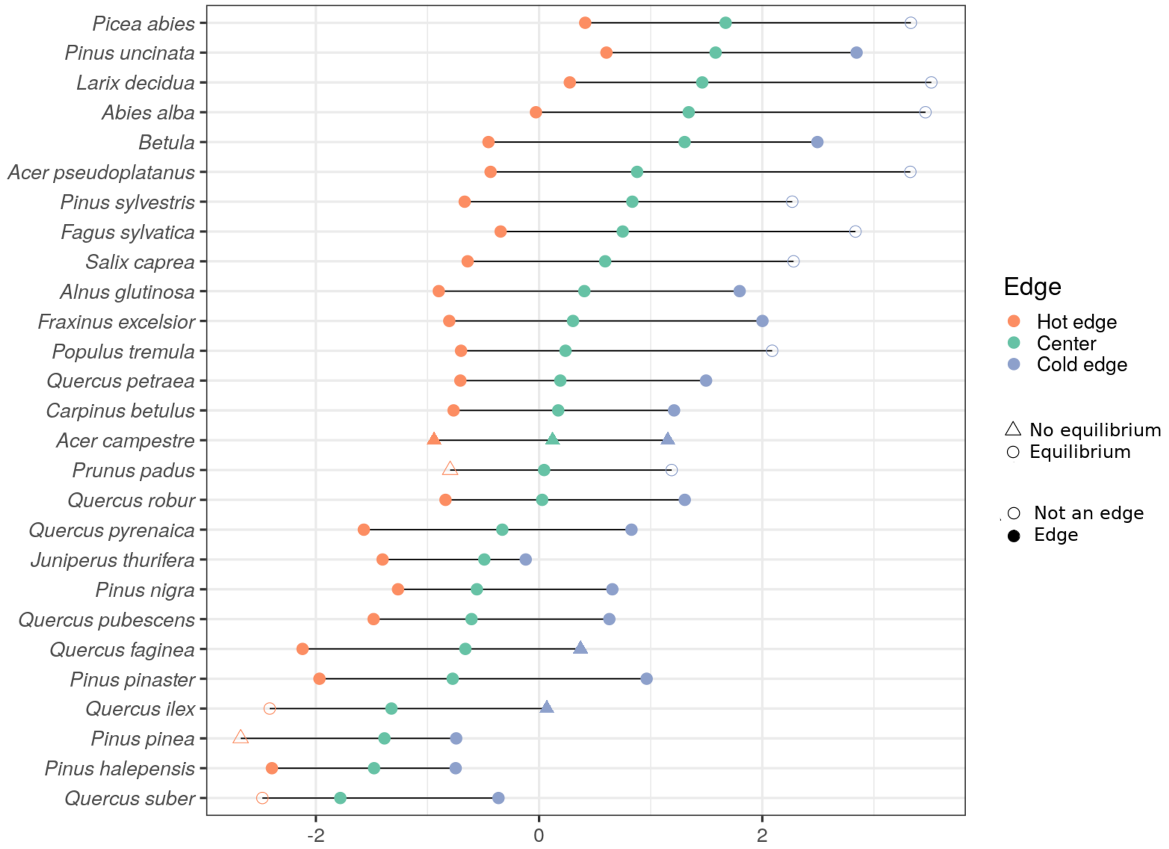

Figure S14: **Species distribution along the first axis of the PCA of the two climatic variables** ***sgdd*** **and** ***wai***. The median of the species distribution along this axis is represented by a green circle and the hot and dry edge and the cold and wet edge by red and blue circle, respectively. Filled circles represent edges selected for the analysis, corresponding to edges where the species distribution models predicted at least a 10% drop in the probability of presence of the species (see above) relative to the distribution center. Point symbols represent climatic conditions where the IPM models resulted in an equilibrium, whereas triangles represent situations where the IPM models did not result in equilibrium and were not analysed.

### 4 Species by species results for $\Omega$

#### 4.1 Recruitment response at the edge

Figure S15 presents $\Omega$ for recruitment at the hot and dry edge and at the cold and wet edge. Figure S16 presents the variation of $\Omega$ with species climatic center.

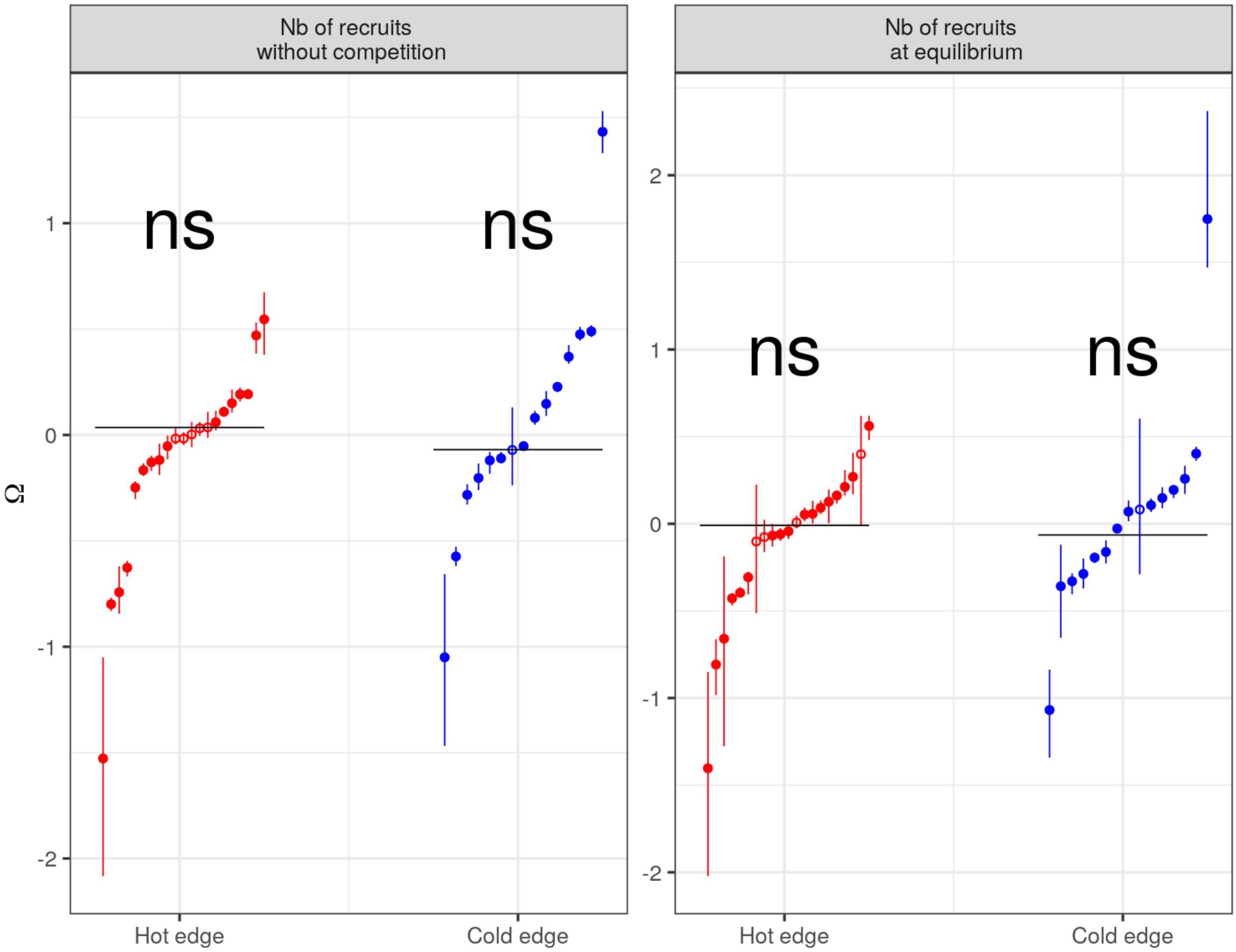

Figure S15: **Relative metrics Ω at the two edges for recruitment rate at low basal area** (left, $1m^2ha^{-1}$) and at the basal area at equilibrium (right).

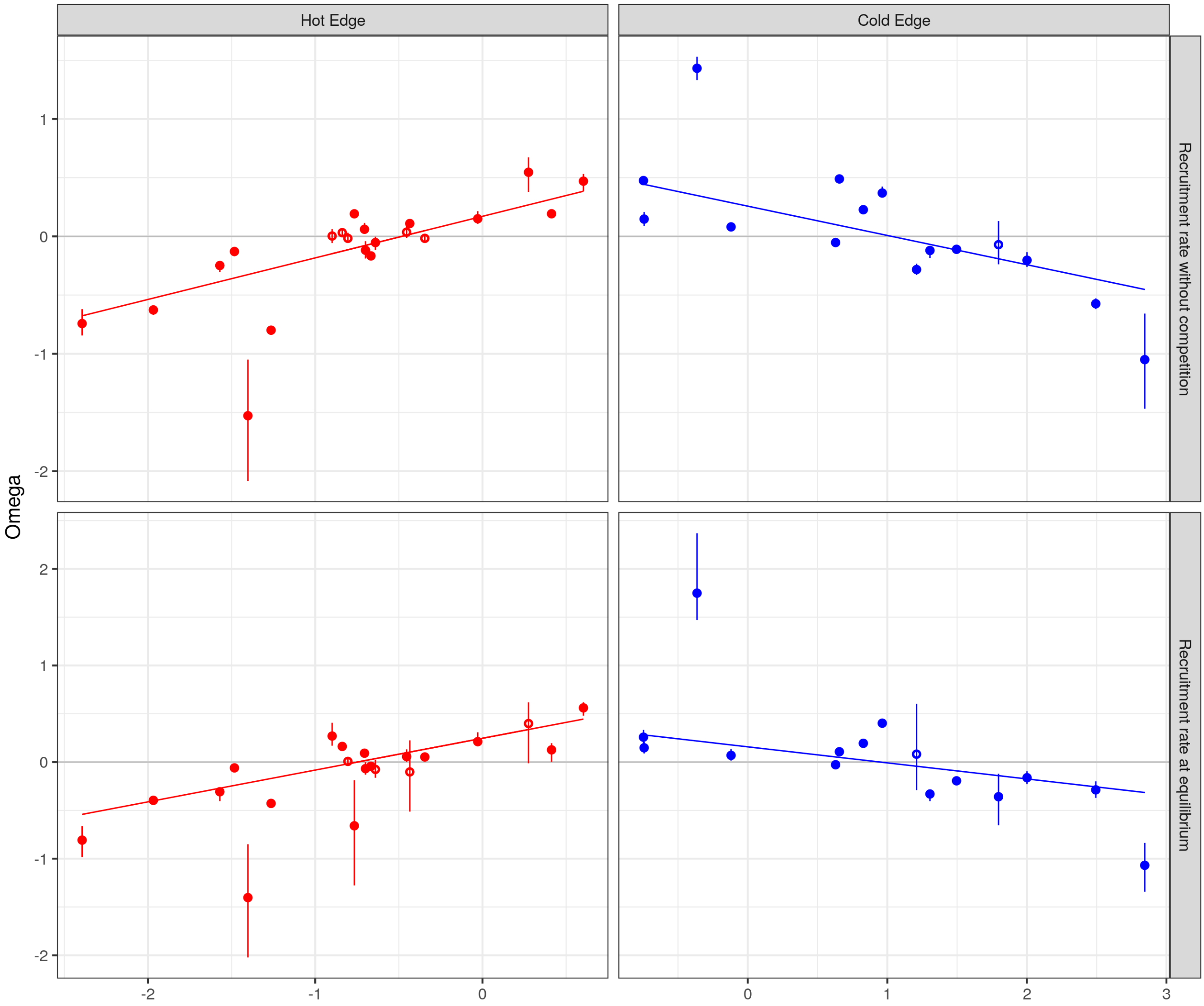

Figure S16: **Relative metrics Ω at the two edges for recruitment rate at low basal area** (left, $1m^2ha^{-1}$) and at the basal area at equilibrium (right).

#### 4.2 Ω by species

Table S7 presents the number of species with Ω significantly different from zero (positive or negative). The mean Ω values for each species and edge type with their associated confidence intervals are accessible in SI file 1 as a ".csv" file. The values of the predicted metrics for the 100 resampled models are provided for each species in a R cran object in the Zenodo repository https://zenodo.org/record/6424490 (file 'outputSimu/Metrics.Rds').

| | Hot Edge<br>22 species | | Cold Edge<br>15 species | |
|---|---|---|---|---|
| Type | Negative | Positive | Negative | Positive |
| Invasion rate at low competition | 12 (55%) | 5 (23%) | 6 (40%) | 7 (47%) |
| Invasion rate at high competition | 14 (64%) | 2 (9%) | 3 (20%) | 8 (53%) |
| Tree density at equilibrium | 11 (50%) | 4 (18%) | 3 (20%) | 9 (60%) |
| Demographic variance | 4 (18%) | 7 (32%) | 6 (40%) | 5 (33%) |
| Time to extinction | 10 (45%) | 1 (5%) | 5 (33%) | 8 (53%) |
| Damping time | 6 (27%) | 4 (18%) | 2 (13%) | 8 (53%) |
| $T^0_{Pert}$ | 5 (23%) | 6 (27%) | 4 (27%) | 7 (47%) |
| $T^{1/2}_{Pert}$ | 6 (27%) | 4 (18%) | 1 (7%) | 4 (27%) |

Table S7: **Number of species with $\Omega$ significantly different from zero (whether negative or positive) for each edge and metric**.

### 5 Phylogenetic regression along climate

We tested the impact of phylogenetic correction on the regression between $\Omega_{sp,i}^{Edge}$ and the median climate of the species. For each edge type (hot and dry vs cold and wet) and for each metric, we performed a Phylogenetic generalised least squares regression (PGLS, see Symonds & Blomberg, 2014) using a phylogeny extracted from Zanne et al. (2014). The phylogenetic tree is presented in Figure S17. Results are summarised in Table S8.

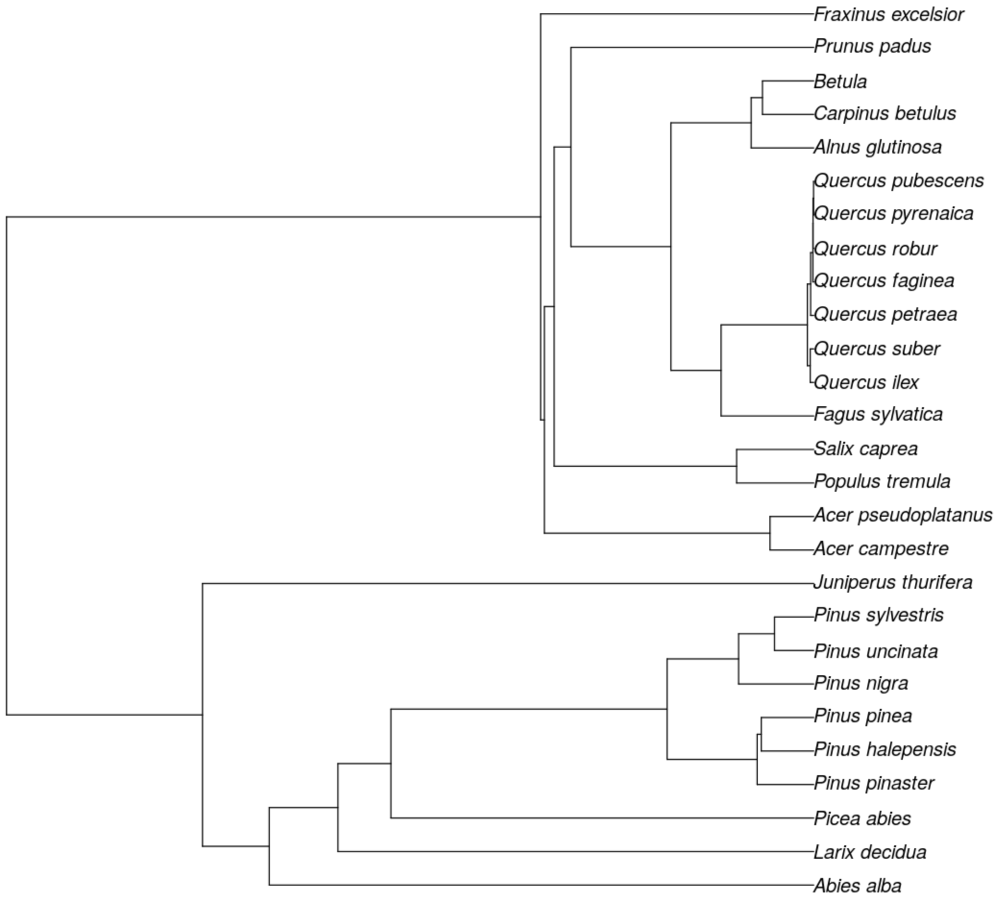

Figure S17: **Phylogenetic tree for species retained in the study and extracted from Zanne et al. (2014).**

| | Hot Edge | | Cold Edge | |
|---|---|---|---|---|
| | **without phylogenetic regression** | **with phylogenetic regression** | **without phylogenetic regression** | **with phylogenetic regression** |
| Invasion rate at low competition | * | * | * | n.s. |
| Invasion rate at high competition | n.s. | n.s. | n.s. | * |
| Tree density | * | * | n.s. | n.s. |
| Demographic variance | * | * | * | * |
| Time to extinction | n.s. | n.s. | n.s. | n.s. |
| Damping time | n.s. | * | n.s. | n.s. |
| Time to zero | * | * | n.s. | n.s. |
| Time to half | * | * | n.s. | n.s. |

Table S8: **Significance of regression between $\Omega$ and climatic conditions in the niche center**. n.s. stands for non significant, * for significant at 5 %.
